# Distinct subpopulations of DN1 thymocytes exhibit preferential γδ T lineage potential

**DOI:** 10.1101/2022.02.25.481936

**Authors:** Seungyoul Oh, Xin Liu, Sara Tomei, Mengxiao Luo, Jarrod P. Skinner, Stuart P. Berzins, Shalin H. Naik, Daniel H.D. Gray, Mark M.W. Chong

## Abstract

The αβ and γδ T cell lineages are both are thought to differentiate in the thymus from common uncommitted progenitors. The earliest stage of T cell development is known as known as CD4^-^CD8^-^ double negative 1 (DN1). These thymocytes have previously been revealed to be a heterogenous mixture of cells, which of only the CD117^+^ fraction have been proposed to be true T cell progenitors that progress to the DN2 and DN3 thymocyte stages, at which point the development of the αβ and γδ T cell lineages diverge. However, recently, it has been shown that at least some γδ T cells are actually derived from a subset of CD117^-^ DN thymocytes. Along with other ambiguities, this suggests that T cell development may not be as straightforward as previously thought. To better understand early T cell development, particularly the heterogeneity of DN1 thymocytes, we performed single cell RNA sequence (scRNAseq) of mouse DN and γδ thymocytes and show that the various DN stages are indeed comprised of transcriptionally diverse subpopulations of cells. We also show that multiple subpopulations of DN1 thymocytes exhibit preferential development towards the γδ lineage. Furthermore, specific γδ-primed DN1 subpopulations preferentially develop into IL-17 or IFNγ-producing γδ T cells. We show that DN1 subpopulations that only give rise to IL-17-producing γδ T cells already express many of the transcription factors associated with type 17 immune cell differentiation, while the DN1 subpopulations that can give rise to IFNγ-producing γδ T cell already express transcription factors associated with type 1 immune cell differentiation.

## Introduction

It is thought that αβ and γδ T cells both arise from common progenitors that seed the thymus from the bone marrow (1, 2). In mice, the earliest stages of T cell development are termed double-negative (DN) because they lack CD4 and CD8 expression. These are further subdivided into DN1 (CD44^+^CD25^-^), DN2 (CD44^+^CD25^+^), DN3 (CD44^-^CD25^+^) and DN4 (CD44^-^CD25^-^) stages. DN1 thymocytes are known to be a heterogenous mixture of cells, based expression of cell surface markers, such as CD24 and CD117 (c-Kit), proliferative capacity and expression of early T lineage genes (3). Of these, the CD117^+^ fraction, which are referred to as “early thymic progenitors” (ETPs), is able to differentiate into DN2 stage cells (3) and appear to be the most efficient at generating CD4^+^CD8^+^ double positive (DP) αβ lineage thymocytes (4). These ETPs are derived from bone marrow progenitors (5), and they still have the capacity to differentiate into NK cells (3, 6, 7) suggesting they remain uncommitted to the T cell lineage. These cells express transcriptional regulators that are associated with stemness as well early regulators of T cell identity (8). ETPs can be further divided into CD24^-^ and CD24^lo^ subpopulations, termed DN1a and DN1b cells, respectively, which are thought to have a precursor-progeny relationship (3). Interestingly, within these ETPs, there is also a CD63^+^Ly6c^+^ subpopulation that appears to be a granulocyte-committed precursor with no T cell potential (8). Thus, even within ETPs, there is significant heterogeneity.

The γδ lineage has been proposed to bifurcate from the main developmental pathway at the DN2>DN3 transition, when *Tcrb/g/d* gene rearrangement occurs (9-12), because clonal assays of ETPs and DN2 thymocytes show that a large proportion at these cells can give rise to both lineages, but this biopotential is lost by the DN3 stage (10). Because thymocytes don’t express cell surface TCR until late DN3, it suggests that the αβ versus γδ commitment occurs independently of a TCR signal. There remains, however, some debate because strong TCR signals can divert TCRβ-pTα (pre-TCR) expressing DN3 thymocytes towards the γδ lineage (13), thus suggesting that there remains some plasticity at the DN3 stage. Subsequently, another study showed that when DN3 thymocytes express both a pre-TCR and γδTCR, the pre-TCR actually contributes to generating a strong TCR signal that drives γδ differentiation (14). Thus, rather than revealing plasticity, strong TCR signaling at the DN3 stage may actually be re-enforcing the differentiation of γδ-committed cells.

Thus, the current favored model of early T cell development is that the ETP subpopulation of DN1 thymocytes is the most immature population in the thymus, which progress to the DN2 stage, at which point commitment toward either the αβ and γδ lineages is initiated. Completion of lineage commitment then occurs at the DN3 stage. The DN3 stage is also when cells are selected for expression of a functional TCRβ, complexed with pre-Tα dimer, which is known as β-selection, or for expression of a complete γδTCR dimer (15).

However, *in vitro* clonal assays have shown that DN3 thymocytes can give rise to a far greater frequency of γδ cells than when starting from DN2 thymocytes (10). This is inconsistent with a simple linear progression model, although this discrepancy could reflect differing survival or plating efficiencies. Moreover, a significant oversight of this model is that it ignores the large number of CD117^-/lo^ DN1 thymocytes that exist. These remaining CD117^-^ DN1 thymocytes can be separated into CD117^lo^CD24^hi^ (DN1c), CD117^-^CD24^hi^ (DN1d) and CD117^-^CD24^-^ (DN1e) subpopulations. However, they were not previously considered part of the T cell developmental pathway because they don’t expand much in culture when compared to the DN1a and DN1b subpopulations (3). Moreover, they appear to differentiate faster than DN2 thymocytes when cultured on OP9-DL1 monolayers, whereas DN1a and DN2b thymocytes progress with a kinetics consistent of being developmentally earlier than DN2 (3).

Potentially, these CD117^-/lo^ DN1 thymocytes may primarily be the progenitors of non-T lineages as it has been shown that dendritic cells can differentiate from these cells as well as from ETPs (16, 17). However, it was recently shown that a subset of IL-17-producing γδ T cells are actually derived from Sox13-expressing DN1d thymocytes, not from ETPs (18), and therefore not via the canonical ETP>DN2>DN3 pathway.

This finding that IL-17-producing γδ T cells can differentiate from DN1d thymocytes also points to another feature of γδ T development that differs from the development of most αβ T cells, that effector outcomes appear to be determined in the thymus rather than in the periphery. First, the expression of IL-17A or IFNγ by mature γδ T cells correlates with Vγ chain usage. Vγ2^+^ γδ T cells tend to produce IL-17A while Vγ1^+^ γδ T cells tend to produce IFNγ (19, 20). Additionally, weak TCR signaling may promote an IL-17A phenotype in γδ T cells, whereas strong signaling promotes an IFNγ phenotype (21).

While there is little controversy to αβ T cell development progressing via the DN2 and DN3 stages and that at least some γδ cells bifurcate at the DN3 stage, the ambiguities described suggests that the αβ versus γδ decision lineage decision is not as strict as the prevailing model and that there is still much to be clarified at these early stages of T cell development. Notably, CD117^-/lo^ DN1 thymocytes cannot be simply omitted from a model of T cell development as highlighted by the finding that at least some IL-17-producing γδ T cells develop from DN1d thymocytes. To better characterize the earliest stage of T cell development, particularly the composition of the DN1 population, we performed single cell RNA sequence (scRNAseq) of mouse DN and γδ thymocytes to determine the transcriptional heterogeneity at single-cell resolution. By delineating transcriptionally distinct subpopulations and assessing their lineage potential we better clarify the composition of the DN populations, particularly of DN1 thymocytes.

## Materials and Methods

### Mice and thymocyte preparations

Thymuses was harvested from C57BL/6 mice at 6 to 7 weeks of age. All experiments were approved by the St Vincent’s Hospital Animal Ethics Committee and performed under the Australian code for the care and use of animals for scientific purposes.

Total thymocytes were obtained by crushing the thymus through a metal sieve to generate a single cell suspension. The cells were washed with PBS and filtered through a 70 *μ*m sieve to remove any clumps. CD4^+^ and CD8^+^ expressing thymocytes were depleted using anti-CD4 and CD8 magnetic-activated cell-sorting (MACS) beads (Miltenyi Biotec), according to the manufacturer’s instructions. The depleted thymocyte preparation was stained with surface antibodies for sorting on a FACSAria (BD Biosciences).

### Flow cytometry

For the analysis of cell surface phenotype, the cells were simply stained with antibodies. For the analysis of intracellular cytokine expression, cells were first restimulated *in vitro* with 50ng/mL phorbol 12-myristate 13-acetate (PMA) + 2 *μ*g/mL ionomycin (both Sigma-Aldrich) in the presence of Monensin (BD Biosciences) at 37°C for 2.5h before staining with cell surface antibodies. The cells were then fixed with the Intracellular Fixation and Permeabilization buffer set (eBioscience) and stained with antibodies against cytokines. All antibodies were purchased from eBiosciences or BD Biosciences. Flow cytometry data were then acquired on a LSR Fortessa III (BD Biosciences) and analyzed with FlowJo software v10.7.0 (Treestar). When only analyzing cell surface phenotype, dead cells were excluded using DAPI. For intercellular cytokine analyses the cells were not stained with DAPI but were gated on live cells determined by size.

### Single-cell RNA sequencing

Sorted thymocytes were counted and checked for viability, then loaded onto the Chromium platform (10X Genomics) for scRNAseq library construction using the Single Cell V2 or V3.1 Reagent Kit according to the manufacturer’s instructions. Libraries were sequenced using 150-cycle/150bp-reads NextSeq500 (Illumina) or 300-cycle/150bp-reads Novaseq (Illumina). Sequencing files were demultiplexed and aligned to *Mus musculus* transcriptome reference (mm10), and count matrix was extracted using CellRanger Single Cell software v2.1.1 or v4.0.0 (10X Genomics) (22).

The Illumina sequencing output was pre-processed with Seurat (v2.3 or v.3.2.2) on R (v3.6.3 and 4.1.0). Cells with <500 genes, >5000 genes or >7% mitochondria gene expression were filtered out as low-quality cells. Following normalization and removal of confounders, highly variable genes were identified and selected using VST selection method (23). Unsupervised linear principal component analysis (PCA) was performed on these highly variable genes to grouped them into 20 principal components. Cell clustering was implemented using the number of components that retain >90% of variance of gene expression in the data.

DoubletFinder (v2.0.3) was applied to remove likely sequencing doublets before downstream analyses (24). The expected number of doublets was calculated as 0.75% of recovered cells. The remaining cells were re-clustered and visualized with t-SNE or uniform manifold approximation and projection (UMAP) dimensional reduction. Differential expression between subclusters was carried out using FindAllMarkers function, with default parameters; differentially expressed genes with adjusted *p*-value <0.05 and fold change >0.5 or <-0.5 (log2FC) were considered unless otherwise stated. The heatmaps was generated using the function DoHeatmap. Hierarchical clustering heatmaps were generated with the R package ComplexHeatmap (v2.8.0) using Euclidean distance measures (25).

Cell cycle genes specific to either the G1, S, G2/M stages was used to perform cell cycle scoring and assign cells to their respective stage of the cell cycle (26). Cell cycle genes were regressed out using Seurat’s built-in regression model.

### Merging of multiple scRNAseq datasets

To compare cell types and proportions across three independent sequencing runs, the datasets were integrated as described at https://satijalab.org/seurat/archive/v3.0/integration.html (27). The Seurat package (v.3.2.2) was used to assemble multiple distinct scRNAseq datasets into an integrated dataset and cell cycling scores were calculated. To remove technical variability, the datasets were pre-processed and normalized using SCTransform (28). To correct for experimental batch effect, integration anchors were identified between the experiments then merged using canonical correlation analysis. Linear dimensional reduction was applied and principal components that retain >90% of variance of gene expression in the data were included for downstream analysis. Unsupervised clustering was implemented on the integrated data.

### Pseudotime trajectory construction

Filtered 10X data was imported into Monocle 2 by generating a cell dataset from the raw counts slot of the Seurat object. Cells were ordered into a branch pseudotime trajectory according to the procedure recommended in the Monocle 2 documentation (29). The highly variable genes identified by Seurat were chosen as ordering genes to recover pseudospatial trajectories using the setOrderingFilter, reduceDimension and orderCells functions in Monocle 2 with default parameters. Differential expression between pseudotime states were determined using the Seurat function FindAllMarkers.

### OP9-DL1 co-cultures

Thymocyte subpopulations of interest were purified by MACS depletion and cell sorting then plated onto OP9-DL1 monolayers (30). The OP9-DL1 cells were inactivate with Mitomycin C (Stem Cell) immediately prior to use. 10^3^ sorted thymocytes were seeded per well in 96-well plates in αMEM (Thermo Fisher) supplemented with 20% FCS (Bovogen Biologicals), penicillin/streptomycin/gentamycin (Sigma), 2ng/mL murine IL-7 (Peprotech) and 5ng/mL human FLT3L (Peprotech). The media was refreshed every 2d and freshly inactivated OP9-DL1 cells were added every 4d.

### Barcode transduction

Sorted total DN1 thymocytes were pre-cultured on OP9-DL1 for 24h. The cells were then transduced with barcode lentivirus library (31) in StemSpan medium (Stem Cell Technologies) by centrifugation at 900 ×g for 1.5h at room temp. A viral titer pre-determined to give 5-10% transduction efficiency was used to ensure that the cells are not transduced with multiple barcodes. 2ng/mL murine IL-7 and 5ng/mL human FLT3L was then added to each well and the cells were returned to the incubator. The following day, fresh αMEM with supplements was added. αβ (CD90.2^+^ CD8α^+^ TCRγδ^-^) and γδ (CD90.2^+^ CD8α^-^ TCRγδ^+^) lineage cells were sorted after 14d and 20d of the OP9-DL1 co-culture.

### Barcode amplification and sequencing

Barcode library construction was performed as described previously (31). The cells were lysed in 0.5mg/ml Proteinase K (Invitrogen) in Direct PCR Lysis Buffer (Viagen) at 55° C for 2h. The Proteinase K was then inactivated at 85°C for 30min and 95° C for 5min. The lysate was split into 2 wells for technical replicate PCRs. A first round of PCR was performed using 1× Standard-Taq magnesium free reaction buffer pack (NEB) with 2 mM MgCl2 (NEB), 0.2mM dNTPs (made in house), 0.5*μ*M TopLiB forward primer (TGCTGCCGTCAACTAGAACA) and 0.5*μ*M of BotLiB reverse primer (GATCTCGAATCAGGCGCTTA) for 32 cycles (1 cycle at 95 °C for 5min, 30 cycles at 95°C for 15sec, 57.2°C for 15sec and 72°C for 15sec followed by 1 cycle at 72°C for 10min). A second round of PCR was then performed to add different Illumina index to each sample by amplifying the first round PCR product with a sample specific Illumina forward index primer and a common Illumina reverse index primer for 32 cycles (1 cycle at 95°C for 5min, 30 cycles at 95°C for 15sec, 57.2°C for 15sec and 72°C for 15sec followed by 1 cycle at 72°C for 10min). An aliquot of the PCR product was run on a 2% agarose gel to check for barcode amplification, then the samples were pooled and the DNA was cleaned using NucleoMag SPRI beads (Machery-Nagel). A 75-cycle sequencing run was performed on a NextSeq instrument (Illumina).

### Cellular barcode data processing and analysis

The data was demultiplexed and aligned to the reference barcode library using *processAmplicons* function from edgeR package (32). The barcode counts were then processed in the following steps: 1) barcodes with <2 read counts in all sample/cell types were excluded from the analysis; 2) Barcodes that were detected in the water control were also removed from the analysis; 3) The read count between technical replicates of the same sample was averaged and the total read count in each sample was then normalized to 100%.

t-distributed stochastic neighbor embedding (t-SNE) was used to the cluster the normalized barcode profiles and visualize any lineage biases of individual barcodes. DBSCAN (version 1.1-8) was then used to classify barcodes based on their corresponding t-SNE coordinates. Heatmaps were plotted to visualize lineage output of barcodes that were identified by transforming the normalized reads value for each barcode using logarithmic transformation. Two-tailed Pearson’s correlation analysis with 95% confidence interval was calculated to determine the correlation between each lineage outcomes using Prism v9 (GraphPad).

### Fetal thymic organ culture (FTOC)

Fetal thymic lobes were isolated from embryos at gestational 14d following timed pregnancies of C57BL/6 female mice. They were cultured for 5-6d on 0.8mm isopore membranes (Millipore) atop surgical gelfoam sponge (Ferrosan Medical Devices) soaked in RPMI-1640 (Sigma) supplemented with 10% FCS (Bovogen Biologicals), 20mM HEPES (Sigma-Aldrich), 50mM 2-mercaptoethanol (Sigma) and 1.35mM 2’-deoxygyanosine (dGuo, Sigma) to deplete endogenous thymocytes. The depleted thymic lobes were then transferred onto new sponges soaked in fresh media with supplements but without dGuo for 2d before repopulation with thymocyte progenitors. To repopulate thymic lobes, they were placed in 20mL hanging drop cultures on Terasaki plates (Sigma-Aldrich) containing 5×10^2^ to 2×10^3^ sorted thymocytes for 24h before returning to fresh sponges. The media was refreshed every 3-4d. Single cell suspensions of the thymic lobes were generated by passing through a 70mm sieve for analysis by flow cytometry.

### Statistical analysis

Statistical testing was performed with one-way analyses of variance (ANOVA) using Prism v9 (GraphPad). *P* values are shown as * < 0.05, ** < 0.01, *** < 0.001, and **** < 0.0001 where each statistical significance was found, and all data are represented as means ± S.E.M.

### Data availability

All scRNAseq datasets have been deposited in the NCBI Gene Expression Omnibus repository under accession number GSE188913.

## Results

### A comprehensive transcriptional map of early T cell development at single-cell resolution

To better characterize the heterogeneity of the early stages of T cell development, we analyzed the transcriptional landscape of DN and γδ thymocytes at single-cell resolution. Cells were sorted from the thymus of C57BL/6 mice for analysis by 10X scRNAseq over three independent runs (Figure 1A and B and Table 1). The first run consisted of total DN (defined as CD4^-^CD8^-^B220^-^CD11b^-^CD11c^-^NK1.1^-^TCRβ^-^) and TCRγδ^+^ thymocytes, the second consisted only of DN1 and DN2 (CD4^-^CD8^-^CD44^+^B220^-^CD11b^-^CD11c^-^NK1.1^-^TCRβ^-^ TCRγδ^-^) cells and the third involved sorting DN1+DN2, DN3 and TCRγδ^+^ cells separately and mixing back together post-sort at a ratio of 55% to 30% to 15%, respectively. This was to ensure that a sufficient number of DN1 and DN2 cells were captured for high-resolution analysis that is not possible by analyzing total DN cells.

**Table 1.** Summary of single-cell RNA sequencing runs

| Parameter | 1<sup>st</sup> 10X run | 2<sup>nd</sup> 10X run | 3<sup>rd</sup> 10X run |
| --- | --- | --- | --- |
| Cell sorting | DN + $\gamma\delta$ | DN1 + DN2 | DN1 & DN2 (55%)<br>$\gamma\delta$ (15%)<br>DN3 (30%) |
| Number of cells loaded | 17,500 | 18,000 | 17,500 |
| Targeted number of cells (recovery) | 10,000 | 10,000 | 10,000 |
| Estimated number of cells (From Cell Ranger) | 5,527 | 8,887 | 8,869 |
| Mean Reads per cell | 82,999 | 58,813 | 45,036 |
| Median Genes per cell | 2,149 | 1,814 | 2,735 |
| Median UMI Counts | 5,982 | 4,830 | 8,343 |
| Filtering (no. cells removed) | 5,253 (274) | 8,851 (36) | 7,990 (879) |
| Detecting highly variable genes | 2,036 HVGs (Feature Selection) | 2000 HVGs (VST Selection) | 2000 HVGs (VST Selection) |
| Dimensionality reduction | 15PCs (90% variance) | 11PCs (91% variance) | 11PCs (90% variance) |
| DoubletFinder (PC distance matrix) |  | pK = 0.005 | pK = 0.24 |
| Number of doublets | 700 | 750 | 750 |
| Doublets removal (Dimensionality reduction) | 2,066 HVGs<br>15PCs (90% variance) |  | 10PCs (90% variance) |
| Resolution (no. clusters) | 2.0 (19) | 2.0 (26)<br>(25- doublets removed) | 2.0 (27) |
| Cell cycle regression (re-clustering) | 14PCs (>90% variance) | 14PCs (>90% variance) |  |
| Cell cycle regression (re-clustering) | 2.0 (19) | 2.0 (25) |  |

**FIGURE 1.**
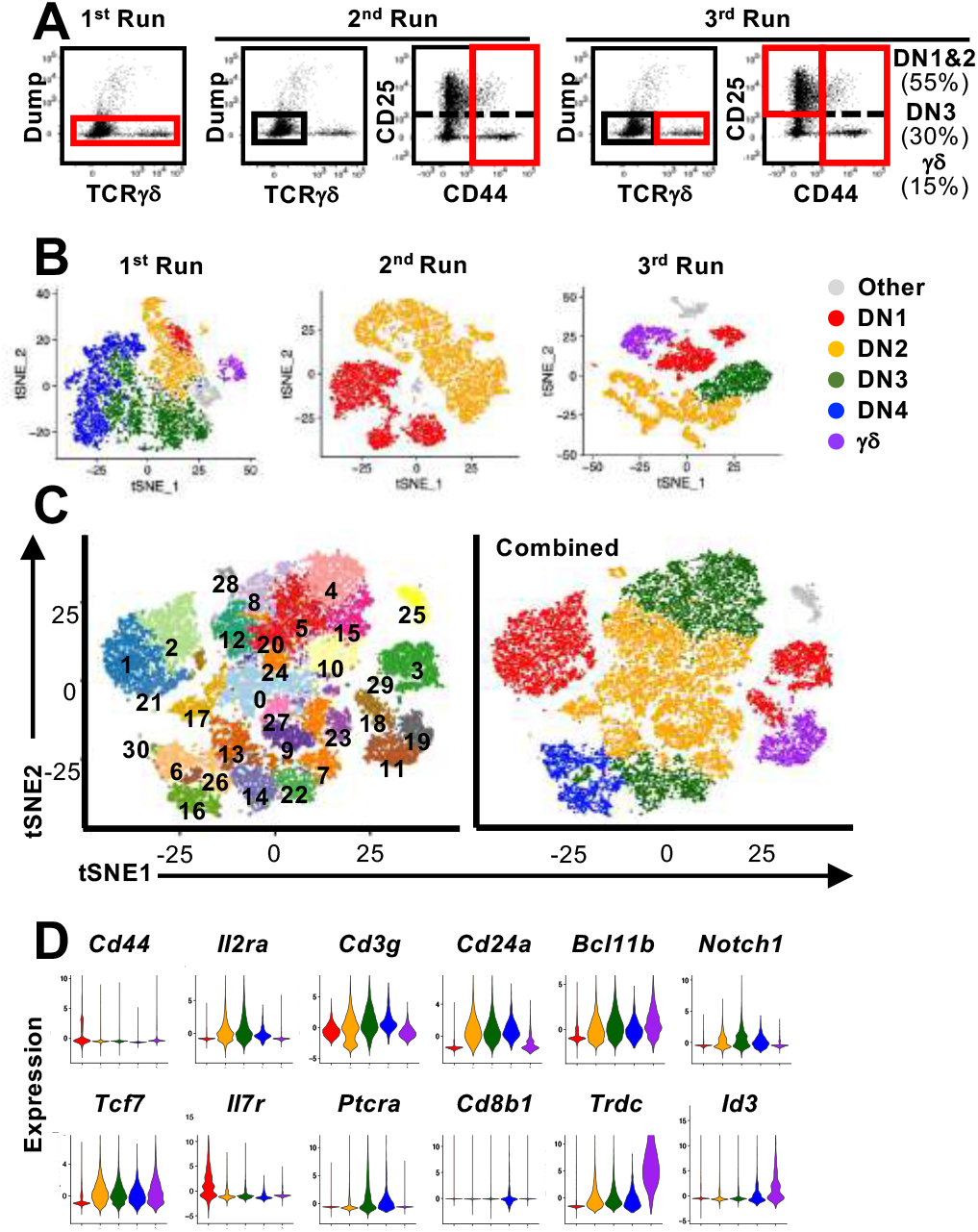
Single-cell RNA sequencing analysis of early T cell development. **(A)** Shown are the gating strategies used to sort DN and γδ thymocytes from C57BL/6 mice for 10X scRNAseq. Three separate runs were completed. The first run was on total DN and TCRγδ^+^ cells. The second run was on only DN1 and DN2 cells. The third run involved sorting DN1 plus DN2, DN3 and TCRγδ^+^ cells separately, which were then mixed back together post-sort at a ratio of 55% to 30% to 15% respectively. Dump = CD4, CD8, B220, CD11b, CD11c, NK1.1 and TCRβ. **(B)** Following processing of the 10X data on CellRanger, each dataset was analyzed for clustering based on the first 12 principal components in Seurat. Shown is the t-SNE plot of each run color-coded to DN developmental stage, TCRγδ^+^ thymocytes or other (non-thymocytes). **(C)** The three datasets, totaling 22,094 cells, were integrated with SCTransform, then clustered with Seurat. A minimal resolution (2.0) was selected such that no cluster contained a mixture of pre- and post-T lineage commitment cells (DN2a versus DN2b) or pre- and post-β-selection cells (DN3a versus DN3b). The resulting clusters (left plot) were then annotated to DN developmental stage, TCRγδ^+^ thymocytes or other (non-thymocytes) (right plot). **(D)** Violin plots showing expression of selected markers used to assign individual clusters to DN development stage.

A total of 22,094 high quality cells passed quality control checks across the three 10X datasets. The three datasets were first integrated by anchoring common cells (27) in order to assemble a global view of early T cell development. To recover biological distinction from the different replicates and minimize batch-associated variability, the pooled data was normalized using SCTransform. Following dimensional reduction, unsupervised clustering was performed using the first 12 PCs. The resolution of clusters was set to a minimum value that was sufficient to separate pre- and post-T lineage commitment cells (DN2a versus DN2b) and pre- and post-β-selection cells (DN3a versus DN3b) into different clusters, which was determined to be 2.0. This identified 30 distinct clusters (Figure 1C), which were assigned to a canonical DN stage or γδ thymocytes based on the expression of key marker genes (Figure 1D and Supplemental Figure 1). High *Cd44* and *Il7r* expression but low T lineage gene expression, including *Il2ra, Tcf7, Cd24a, Notch1 and Bcl11b*, identified DN1 cells. DN2 cells were identified by upregulation of T lineage genes and downregulation of *Il7r*. DN3 cells were identified by *Ptcra* and further upregulation of T lineage genes. Low *Cd8b1* and loss of *Il2ra* distinguished DN4 from DN3 cells. High *Trdc, Id3, Sox13* and *Rorc* identified γδ thymocytes. This indicates that multiple distinct populations correspond to each of the canonical DN stages.

Because there is a massive expansion of cell number between the ETP and DP stages, we also tested the effect of cell cycle gene expression on the clustering. Cell cycling scores were calculated from the integrated data and regressed using Seurat’s built-in regression model (Supplemental Figure 2). There was a slight variation in the number of output clusters, but we did not observe any biological variability that could be explained by cell cycle status. The genetic profiles were nearly identical between the outputs with cell cycle genes left in and regressed out (Supplemental Figure 2). Thus, the transcriptional heterogeneity of DN thymocytes observed is not simply a result of being in different phase of the cell cycle. This also meant that exclusion of cell cycle genes was not necessary for downstream analyses.

**FIGURE 2.**
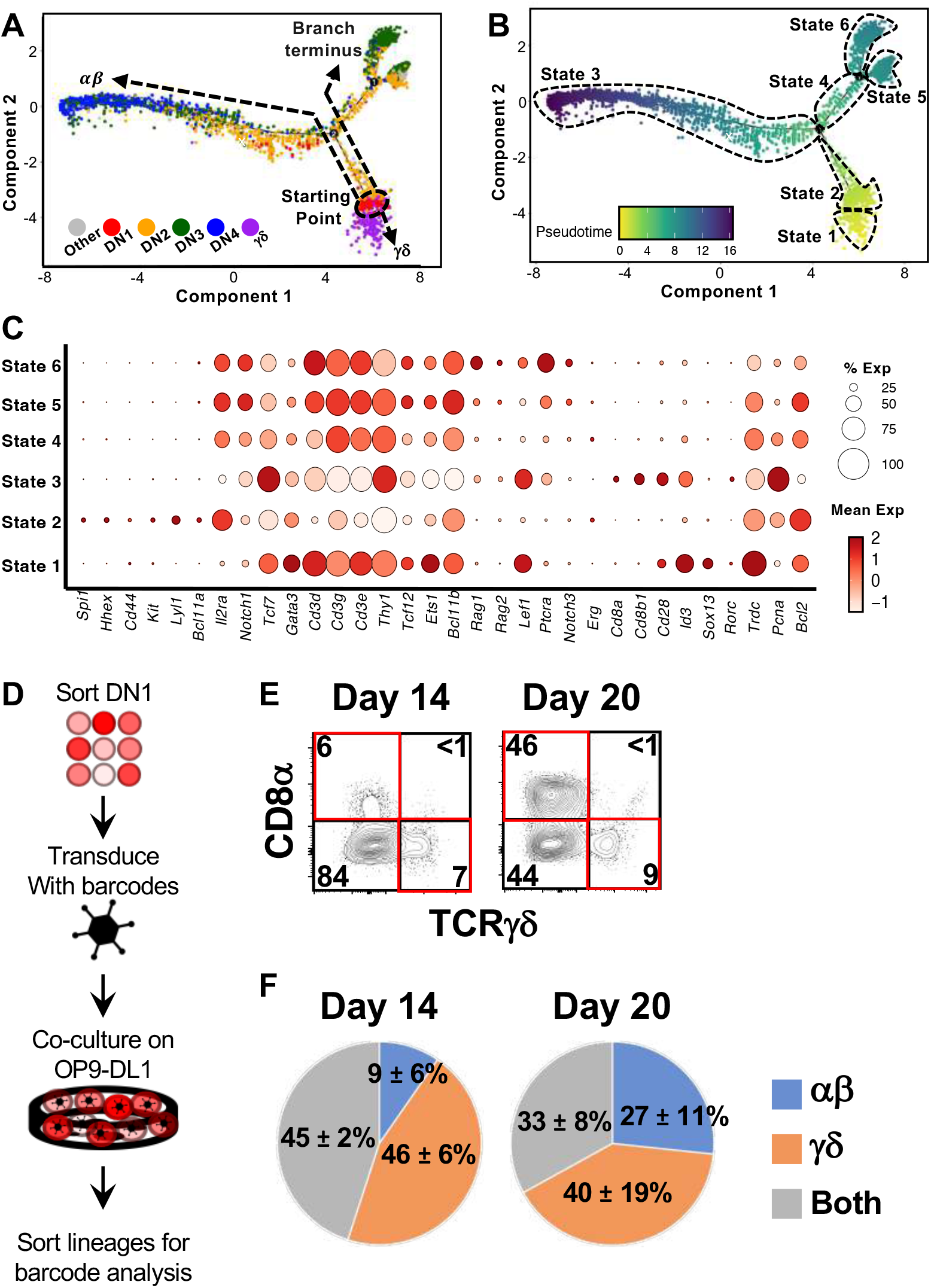
Trajectory analysis suggest that at least some γδ thymocytes develop directly from DN1 thymocytes. **(A)** Pseudotime analysis of total DN and γδ thymocytes (first 10X run) with Monocle 2. The cells are color-coded by thymocyte development stage (DN 1 to 4 or γδ) based on expression of key marker genes (described in Supplemental Figure 1) by the individual clusters. **(B)** Six distinct states were identified within this asymmetric trajectory. **(C)** Dot plots for expression of key genes across the six states. Dot size indicates the percentage of cells with expression, while color saturation indicates average expression level within the state. **(D)** Overview of the experimental setup to track lineage outcomes of DN1 thymocytes differentiated in OP9-DL1 cocultures by cellular barcoding. Total DN1 (CD25^-^ CD44^+^CD4^-^CD8^-^B220^-^CD11b^-^CD11c^-^NK1.1^-^TCRβ^-^TCRγδ^-^) thymocytes were sorted the thymus of C57BL/6 mice and tagged with unique genetic barcodes by transduction with a lentiviral library encoding the barcodes. The cells were then differentiated on OP9-DL1 monolayers. Half the culture was sorted at day 20 for αβ (CD90.2^+^CD8^+^CD4^+/-^TCRγδ^-^) and γδ (CD90.2^+^TCRγδ^+^) lineage cells. The remaining half was differentiated to day 20 and then sorted. **(E)** Example FACS profiles at day 14 and 20 of cultures. **(F)** The barcode composition of the αβ and γδ populations at the two time points were analyzed by Illumina sequencing. Shown is the fraction of unique barcode sequences that were found only in the resulting αβ cells, γδ cells or in both populations for that time point. The values indicate the mean ± S.E.M. of two independent sort/transduction experiments.

### Trajectory analysis implies that γδ T cell development branches from DN1 thymocytes

We next employed Monocle 2 (29) to infer a potential developmental pathway from the scRNAseq data of DN and γδ thymocytes by ordering the cells based on tracking gene expression in pseudotime analysis (Figure 2A). This assembled the cells along an asymmetric trajectory that divided into six states (Figure 2B). Each state was then analyzed for expression of signature genes (Figure 2C) in order to assign a stage in development relative to β-selection. State 2, comprising DN1 and some DN2a cells, was identified as the starting point. State 3 corresponds to the main αβ pathway, with DN4 as the endpoint. States 5 and 6 terminated with DN3 cells and appears to correspond to cells that have not pass β-selection because differential gene expression analysis between these and State 3 found an enrichment of genes associated with “Cellular senescence” and “Apoptosis” KEGG pathways, with adjusted *P* values (-Log_10_) of 6.77 and 6.20, respectively (data not shown). State 1 corresponds to the γδ branch. Thus, based simply on the transcriptomic profile of the cells, Monocle 2 appears to infer that γδ cells develop directly from DN1. We also performed trajectory analyses with another algorithm, Slingshot (33), and it also suggested that γδ cells develop directly from DN1 (not shown). Of course, these are only computational predictions.

### Cellular barcoding in OP9-DL1 cocultures suggests only a partial overlap of the DN1 thymocytes that give rise to αβ and γδ T cells

If the αβ and γδ lineages can develop independently, we might expect to see each lineage being derived from distinct DN1 thymocytes. To investigate this, we sorted total DN1 (CD44^+^CD25^-^CD4^-^CD8^-^B220^-^CD11b^-^ CD11c^-^NK1.1^-^TCRβ^-^TCRγδ) thymocytes, which includes both CD117^+^ ETPs and the CD117^-/lo^ subpopulations. The cells were tagged with a lentiviral library of unique DNA barcodes (31). Once tagged, that barcode is inherited by the progeny of that cell and thus we can estimate how frequently αβ cells and γδ cells originate from the same starting DN1 cell (Figure 2D). If an αβ cell and a γδ cell inherit the same barcode sequence, it means that they were derived from the same progenitor. CD8^+^CD4^+/-^ and TCRγδ^+^ cells were sorted after 14d and 20d of culture and analyzed for barcode composition (Figure 2E). Cell surface CD4/8 was used as the marker of αβ lineage cells because the fixation required to detect intracellular TCRβ interferes with downstream analyses. Late DN4 and DP-staged cells are captured by this sort-strategy. To determine which barcodes propagated from the DN1 thymocytes, the resulting αβ and γδ populations were amplified by PCR then Illumina-sequenced (Figure 2F). We found that at day 14, only 45% of the detected barcode species were sequenced in both αβ and γδ populations, while at day 20, only 33% of detected barcode species were sequenced in both. At both time points, the majority of unique barcode species were detected in only the αβ or γδ population, suggesting that these were derived from DN1 thymocytes that gave rise to only one lineage. Together with the trajectory analysis of the scRNAseq data, this suggest that there are different DN1 thymocyte subpopulations, some of which are biopotential for both lineages, while others appear preferentially developed into either αβ cells or γδ cells. Thus, it could be possible that γδ cells do indeed develop directly from DN1 thymocytes rather than following the canonical pathway and bifurcating from αβ cells at the DN2>DN3 stage.

### Transcriptional heterogeneity of DN1 and DN2 thymocytes

To better delineate the earlier stages of the developmental model, we focused the second 10X run that was performed specifically on DN1 and DN2 thymocytes. Unsupervised clustering of the 8,851 cells that pass quality control checks identified 26 clusters (Figure 3A). This was based on a resolution of 2.0, which was the minimum number that resulted in clusters containing only DN1, DN2a or DN2b thymocytes. Specifically, we checked that no cluster appeared to contain a mixture of cells with DN2a (pre-T-specification) or DN2b (post-T-specification) identity, at least based on expression of conventional stage marker genes. DN1s expressed high levels of progenitor markers, including *Hhex, Il7r* and *Cd44* but low levels of T-commitment markers, such as *Il2ra, Tcf7, Notch1, Cd24a* and *Myb* (Supplemental Figure 3). Late T-lineage commitment genes, including *Bcl11b, Rag1, Rag2, Notch3* and *Ptcra*, were used to separate the DN2a and DN2b subpopulations (Supplemental Figure 3). This resulted in eight DN1, five DN2a and 10 DN2b subpopulations. Some clusters formed distinct subpopulations, particularly DN1 thymocytes, while others appeared to be divisions within a continuum, particularly DN2 thymocytes. Such heterogeneity within these early T cell developmental stages is consistent with that previously reported, at least for DN1 thymocytes (3). There were also three small clusters of non-T cells consisting of doublets and B cells that, for simplicity, were excluded from downstream analysis.

**Figure 3.**
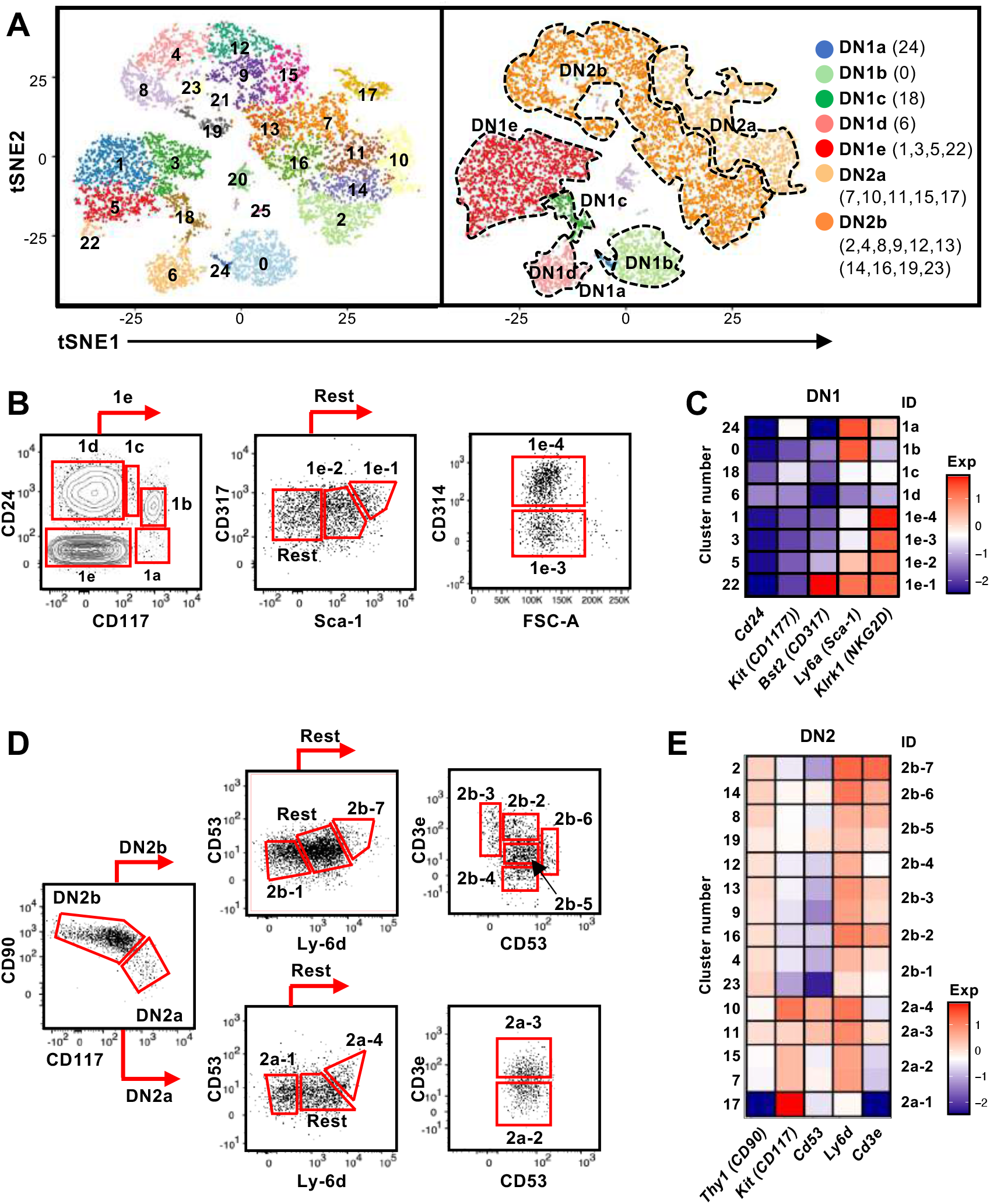
Identification of cell surface markers for delineating the DN1 and DN2 subpopulations inferred from scRNAseq. **(A)** t-SNE visualization of 8,851 DN1 and DN2 thymocytes from the second 10X run clustered with Seurat. A minimal resolution (2.0) was selected such that no cluster contained a mixture of pre- and post-T lineage commitment cells (DN2a versus DN2b). This yielded 26 clusters (left plot), which were then annotated as DN1a, DN1b, DN1c, DN1d, DN1e, DN2a or DN2b (right plot). **(B)** The DN1 subpopulations were analyzed for differentially expressed genes encoding cell surface proteins. Antibodies against these proteins were then tested. Shown is the flow cytometric strategy to subdivide the eight DN1 subpopulations using a panel of five antibodies. Total DN1 cells were identified as CD25^-^ CD44^+^CD4^-^CD8^-^B220^-^CD11b^-^CD11c^-^NK1.1^-^TCRβ^-^TCRγδ^-^. **(C)** Heatmap showing the average expression of the genes encoding cell surface markers used to delineate the DN1 subpopulations. **(D)** The flow cytometric strategy to subdivide four populations of DN2a cells and seven populations of DN2b cells using a panel of five antibodies. Total DN2 cells were identified as CD25^+^CD44^+^CD4^-^CD8^-^B220^-^ CD11b^-^CD11c^-^NK1.1^-^TCRβ^-^TCRγδ^-^. **(E)** Heatmap showing the average expression of the genes encoding the cell surface markers use to delineate the DN2 subpopulations.

### Identification of novel cell surface markers for delineating DN1 and DN2 subpopulations

While various cell surface markers have been used to divide DN1 thymocytes into DN1a to DN1e, and DN2 thymocytes into DN2a and DN2b (3), these are insufficient to delineate the larger number of subpopulations that are suggested by the scRNAseq analysis. We therefore needed to identify useful cell surface markers that could be used for flow cytometry. Differential expression analysis was performed to select features (*P*<0.05 and 2-fold difference) for cross-referencing with GO terms for cell surface proteins (not shown). We then tested commercial antibodies against these candidates. Protein does not always correlate with mRNA levels and therefore not all antibodies produced a staining pattern that matched the scRNAseq data. However, we were able to identify a minimal panel to delineate the eight DN1 subpopulations and most of the DN2 subpopulations.

CD24 and CD117 have previously be shown to divide DN1 thymocytes into five subpopulations (3). DN1a, DN1b, DN1c and DN1d each corresponded to a single cluster in the scRNAseq analysis, but we identified four clusters (#1, 3, 5 and 22) that corresponded to DN1e thymocytes (Figure 3A). The inclusion of CD314 (NKG2D), CD317 (BST2) and Sca-1 delineated these four DN1e subpopulations (Figure 3A and B).

DN2a and DN2b cells can be divided based on expression of several markers including CD90 and CD117, which were then subdivided further based on CD53, Ly-6d and CD3e expression (Figure 3D and E). The pair of DN2a clusters 7/15 could not be separated by cell surface markers as well as the pairs of DN2b clusters 4/23, 9/13 and 8/19 due to very similar gene expression profiles. Thus, these cluster pairs were combined. The resulting 11 DN2 clusters were annotated as DN2a-1 to 4 and DN2b-1 to 7.

### OP9-DL1 cultures reveal that specific subpopulations exhibit biases towards either the γδ lineage

The transcriptional heterogeneity of the DN1 thymocytes may be an indication that only some subpopulations are true progenitors of T cells, as previously suggested (3). Alternately, these different subpopulations could represent different progenitors of αβ and γδ lineages, which would be consistent with the early branch point inferred by pseudotime trajectory analyses (Figure 2A and B). To test this, we sorted the DN1 and DN2 subpopulations using our antibody panels and accessed their αβ versus γδ lineage potential in OP9-DL1 cultures after 14d and 20d. Expression of TCRγδ identified γδ lineage cells. Expression of CD8α identified αβ lineage cells, which captures cells from late DN4 to DP stages and the few CD8 (TCRβ^+^) single positive cells that generated. We also checked CD4 expression (not shown), which correlates entirely with the DP stage. CD4 (TCRβ^+^) single positive cells do not develop in these cultures due to lack class II MHC presentation (34).

At day 14, there were primarily αβ lineage or undifferentiated cells in DN1a, DN1b and DN1c cultures, with very few TCRγδ^+^ cells produced (Figure 4A to C). DN1d and all DN1e cultures only generated TCRγδ^+^ cells. Notably, the number of TCRγδ^+^ cells produced in all DN1e cultures was substantially greater than the number of TCRγδ^+^ produced in DN1a and DN1b cultures (Figure 4C).

**FIGURE 4.**
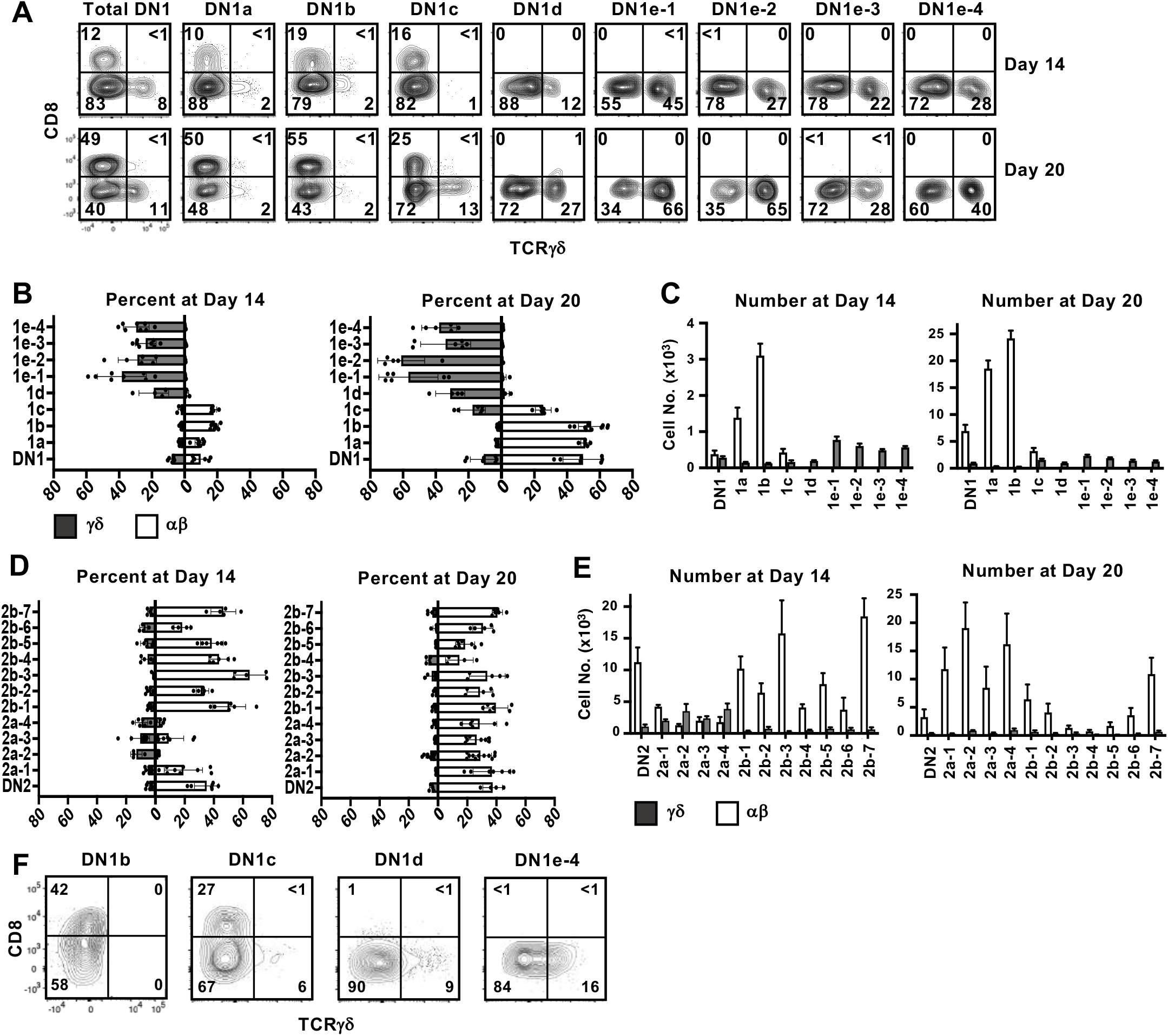
Assessing the αβ or γδ potential of DN1 subpopulations. **(A)** Sorted DN1 subpopulations were cultured on OP9-DL1 monolayers to assess their lineage potential. The cultures were analyzed at 14d and 20d of culture by flow cytometry. αβ lineage cells were identified as CD8^+^ (capturing late DN4 and DP cells) while γδ lineage cells were identified as TCRγδ^+^. Representative flow cytometric plots are shown. (**B, C)** Pooled data analyzing αβ versus γδ differentiation from sorted DN1 subpopulations. The means ± S.E.M. of four to nine replicates performed over four independent experiments are shown. See Supplemental Tables 1 and 2 for *P*-value calculations. **(D, E)** Pooled data analyzing αβ versus γδ differentiation from sorted DN2 subpopulations. The means ± S.E.M. of four to nine replicates performed over four independent experiments are shown. See Supplemental Tables 3 and 4 for *P*-value calculations. **(F)** Repopulation of dGuo-depleted FTOCs with select DN1 subpopulations. The lobes were then analyzed by flow cytometry for CD8 versus TCRγδ expression after 14d. Shown is a representative from three independent experiments.

At 20d, there was a substantial increase in the percentage and number of αβ cells produced in DN1a and DN1b cultures, while DN1c had given rise to both γδ cells and αβ cells. DN1d and DN1e subpopulations continued to exhibit a bias toward the γδ lineage, with DN1e-1 and DN1e-2 producing the highest percentage and number of TCRγδ^+^ cells. In terms of absolute numbers, DN1a and DN1b subpopulations produced αβ cells at a greater rate than the production of γδ cells from the DN1c, DN1d or DN1e subpopulations. Starting from 10^3^ DN1a thymocytes, almost 2×10^4^ αβ cells were present after 20 days, whereas only 2-3×10^3^ γδ cells had been produced from all four DN1e cultures after this time (Figure 4C). This of course could be a reflection of γδ differentiating and dying more quickly, but is also implies that DN1a and DN1b subpopulations are proliferating more quickly (3) and preferentially producing αβ cells.

At day 14, all DN2b subpopulations and DN2a-1 had produced a high percentage and number of αβ cells and few TCRγδ^+^ cells, while the rest of DN2a subpopulations produced mostly TCRγδ^+^ cells and some αβ cells (Figure 4D/E). By day 20, all DN2 subpopulations had produced αβ lineage cells, while TCRγδ^+^ cells had been lost from the DN2a cultures. There was also a dramatic reduction in cell numbers in the DN2b cultures, which may be a result of the cells undergoing cell death after reaching the DP stage because these cultures poorly support the later stages of αβ maturation (34).

### Fetal thymic organs cultures confirm the lineage bias of DN1 subpopulations

While OP9-DL1 cultures are a well-characterized system for analyzing T cell development, it is possible that the lineages biases observed for the different DN1 subpopulations may be exaggerated in this model. We therefore also assessed the differentiation of select DN1 subpopulations in fetal thymic organ cultures (FTOCs), by seeding the sorted subpopulations into dGuo-depleted fetal thymic lobes. Like in the OP9-DL1 co-cultures, DN1b cells preferentially produced αβ cells, DN1c cells primarily produced αβ cells and some γδ cells, while DN1d and DN1e-4 cells only produced TCRγδ^+^ cells (Figure 4F). Thus, both OP9-DL1 and FTOC systems suggest that distinct DN1d and DN1e subpopulations are biased toward the γδ lineage. Moreover, although DN1a, DN1b and DN1c can produce both αβ cells and γδ cells, they are heavily biased toward the αβ lineage and produce these cells in large numbers. The DN1d and DN1e subpopulations, together, make up more than three-quarters of DN1 thymocytes. Thus, on a per cell basis, it appears that more γδ cells are produced from CD117^-^ DN1 thymocytes than from what are traditionally considered the ETPs (DN1a and DN1b).

### Gene expression analysis of DN1d and DN1e subpopulations suggest potential relationships with distinct mature γδ subsets

Consistent with two group of γδ progenitors identified by Sagar and colleagues (35), γδ thymocytes clearly segregate into two clusters (#11 and 9) in our scRNAseq dataset (Figure 1C). We therefore wanted to determine how these relate to the different DN1 subpopulations that produced γδ cells. Differential gene expression analysis revealed substantial differences between the two γδ clusters, with 93 genes expressed at significantly higher levels by cluster 11 cells, while 117 genes were expressed at significantly higher levels by cluster 19 cells (Figure 5A). Genes that were highly expressed by cluster 11 include *Gzma, Blk, Maf, Sox13, Etv5, Gata3, Ccr9, Rorc, Sox4, Tcf12, Lgals9, Cmak4* and *Bcl11b*, which are all associated with the IL-17 effector phenotype (35). This suggests that cluster 11 cells are probably the γδ thymocytes that mature into the γδ17 subset in the periphery. Numerous interferon-related genes, such as *Stat1*, were highly expressed by cluster 19 cells, suggesting that these cells probably mature into the IFNγ-producing γδ subset. This is consistent with a previous study that suggested the eventual effector function of γδ T cells may already be acquired in the thymus (21).

**FIGURE 5.**
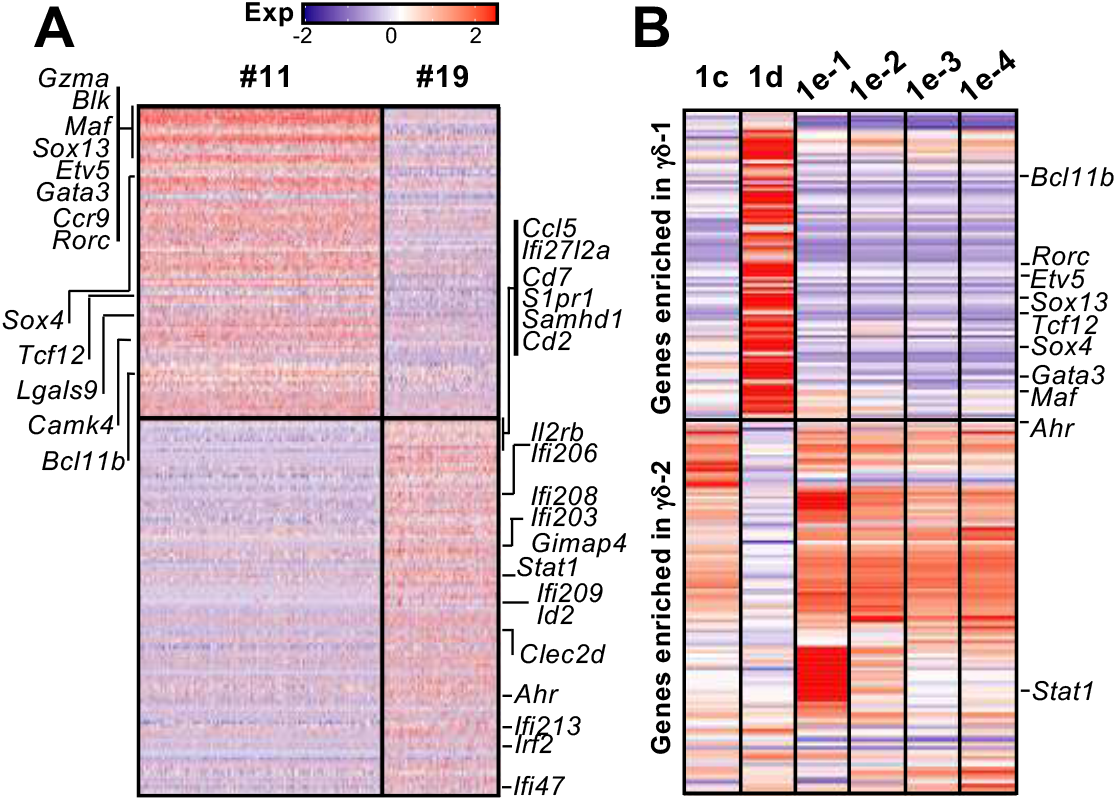
The gene expression profiles of the different DN1 subpopulations correlate with distinct γδ effector subsets. **(A)** Heatmap showing genes differentially expressed (*P*-value <0.05 and 2-fold difference) between the two γδ thymocyte subpopulations (clusters 11 and 19) identified in the scRNAseq analysis of DN and γδ thymocytes in Figure 1C. Each column is an individual cell in the dataset while each row is a differentially expressed gene. Genes associated with either IL-17A (left side) or IFNγ production (right side) are indicated. **(B)** Analysis of DN1 subpopulations for expression of the 210 genes differentially expressed between the two γδ thymocyte populations. Indicated are transcription factors that are associated with either IL-17A or IFNγ-producing effector subsets.

Next, to investigate the relationship between these γδ thymocyte subpopulations and the DN1 subpopulations that differentiate into TCRγδ^+^ cells, the DN1c, DN1d and DN1e subpopulations were analyzed for the expression of the 210 differentially genes that distinguished the two γδ thymocyte populations. This revealed a similar transcriptional profile between DN1d and cluster 11 γδ thymocytes, while DN1c and all the DN1e subpopulations overlapped significantly with cluster 19 γδ thymocytes (Figure 5B). However, there were clearly also differences between the DN1e subpopulations.

We then focused on transcription factors because they are regulators of gene expression and could potentially play key roles in hardwiring γδ effector outcomes in DN1 thymocytes. *Sox13* was highly expressed by DN1d cells (Figure 5B), which was previously shown to be important for the differentiation of a subset of DN1 thymocytes into IL-17-producing γδ T cells (18). Interestingly, DN1e subpopulations also express transcription factors associated with IL-17-producing γδ T cells. Notably, DN1e-1 and DN1e-2 cells expressed high levels of *Maf*, while *Gata3* was highly expressed by both DN1e-1 and DN1d cells (Figure 5B). We thus predict that the foundation of γδ effector transcriptional network may already be in place in DN1 subpopulations, with DN1d going on to develop into IL-17-producing cells and DN1e potentially producing both effector subsets.

### Different DN1 subpopulations can give rise to different γδ effector subsets

To determine if different DN1d and DN1e subpopulations might differentiate into distinct effector γδ T cell subsets, the TCRγδ^+^ cells that developed in OP9-DL1 co-cultures were analyzed for intracellular IL-17A and IFNγ expression and for expression of specific Vγ chains (Fig 7A). Cytokine production is highly associated with specific Vγ chain usage, with Vγ1^+^ cells enriched for IFNγ and Vγ2^+^ cells enriched for IL-17A production (19, 36).

DN1c thymocytes generated both Vγ1^+^ and Vγ2^+^ cells, with only a low percentage of Vγ1^+^ cells expressing IFNγ (Figure 6B to D). DN1d primarily produced Vγ2^+^ cells that express IL-17A. Similarly, DN1e-1 and DN1e-2 were biased towards a Vγ2^+^ IL-17A^+^ γδ effector subset. On the other hand, only DN1e-3 and DN1e-4 exhibited the plasticity to generate both IL-17A and IFNγ-expressing cells (Figure 6B to D). This suggests that the foundation of γδ effector programs are already in place in DN1 thymocytes.

**FIGURE 6.**
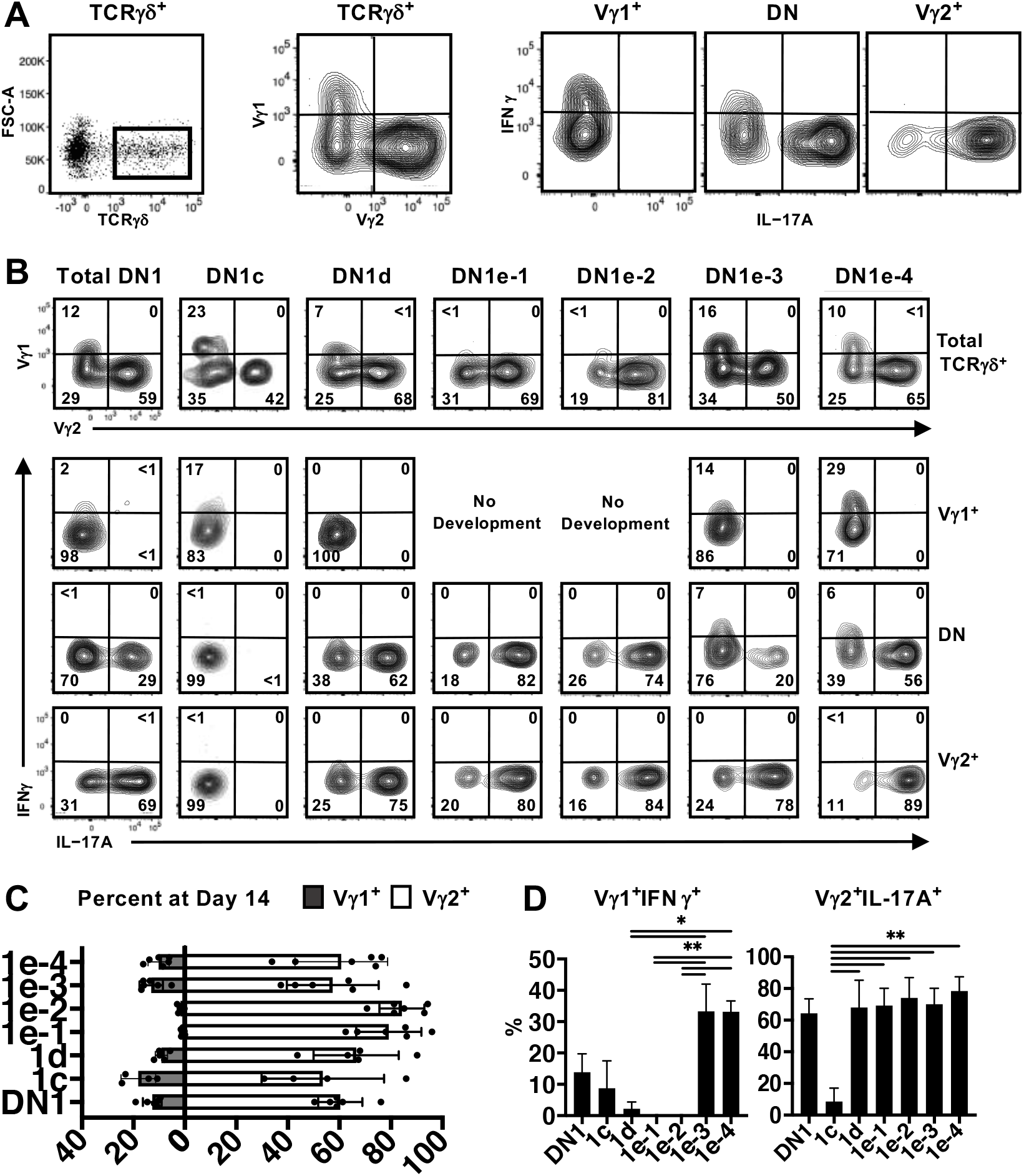
Analysis of γδ effector outcomes from DN1 subpopulations. **(A)** Gating strategy for analyzing the phenotype of γδ T cells generated from DN1 subpopulations after culturing on OP9-DL1 monolayers for 14d. TCRγδ^+^ (CD4^-^CD8^-^TCRβ^-^) cells were first divided based on Vγ1 versus Vγ2 expression. The three subpopulations, including Vγ1^-^Vγ2^-^ double negative (DN) cells were then analyzed for intracellular IL-17A and IFNγ expression. **(B)** Shown are the flow cytometric plots from a representative experiment. The top row shows the Vγ1 versus Vγ2 expression on gated TCRγδ^+^ cells. The Vγ1^+^, Vγ2^+^ and double negative (DN) cells were then analyzed for IL-17A versus IFNγ expression in the bottom three rows. **(C)** Pooled data analyzing the percentage of Vγ1 versus Vγ2 cells differentiated from sorted DN1 subpopulations. The means ± S.E.M of four to six replicates performed over three independent experiments is shown. See Supplemental Table 5 for *P*-value calculations. **(D)** Pooled data analyzing percentage of Vγ1^+^IFNγ^+^ (left) and Vγ2^+^IL-17A^+^ (right) cells out of total TCRγδ^+^ cells. The means ± S.E.M is shown (\**P*<0.05, \*\**P*<0.01).

## Discussion

Our study has confirmed that, at a transcriptional and functional level, DN1 thymocytes are indeed a heterogenous population of cells (3). We also confirmed that DN1a and DN1b thymocytes (CD117^+^ DN1 fraction), which together have been considered the true ETPs, can give rise to both αβ and γδ lineages. However, we showed that γδ cells can be derived from multiple progenitor sources, with the CD117^-^ DN1 subpopulations more efficient at producing γδ cells than DN1a and DN1b subpopulations. That being said, αβ cells are produced in far greater numbers because DN1a and DN1b thymocytes are have a great proliferative capacities compared to the other subpopulations (3). This would explain why TCRγδ^+^ thymocytes are greatly outnumbered by αβ lineage thymocytes (DN4 onwards) within the thymus.

While CD117^+^ DN1 cells have been considered bipotential, a previous study demonstrated the existence of lineage-restricted precursors. Spidale and colleagues showed that neonatal IL-17-producing γδ T cells develop from a subset of Sox13-expressing DN1d thymocytes and that this lineage is determined by a cell intrinsic program that is independent of TCR signaling (18). Our study has now expanded the subdivision of DN1 thymocytes to defined eight subpopulations. We showed that not only are DN1d thymocytes primed towards the γδ lineages but also the four DN1e subpopulations. This does not mean lineage commitment is determined entirely at this point because the progression from DN1 to fully mature γδ thymocytes still requires TCR signaling to select cells that express a functional TCRγδ dimer. Indeed, Scaramuzzino and colleagues showed that TCR signaling is required for γδ maturation because LAT-deficient DN3 cells that express a γδ TCR are unable to completely activate the γδ lineage program, including expression of *Sox13, Maf*, and *Cpa3* (37).

Evidence suggests that TCR signal strength is an important determinant of lineage outcome in bipotential precursors. Strong signals that activate the ERK-EGR-ID3 pathway have been shown to drive γδ T cell differentiation, whereas a weak signal promotes the αβ fate (38-40). However, it is still unclear whether TCR signal strength is dependent on instructive extrinsic signals or simply a result of stochastic selection of the TCR chains that are expressed. The instructive model proposes that TCRγδ signals competes with pre-TCR(β) signals and that the lineage decision is determined by cell specific interactions that activate key transcription factors, which in turn instructs gene expression program (15). In contrast, the stochastic selective model postulates that lineage commitment is determined prior to the onset of TCR gene rearrangement, and it is only when a thymocyte receives the appropriate TCR signal that matches its hardwired identity that it actually progress along the αβ or γδ developmental pathway. DN1d and DN1e thymocytes are clearly committing to γδ lineage cells prior to the expression of the TCR. The fact that when a DN3 thymocyte expresses both a pre-TCR and γδTCR results in an even stronger signal that drives γδ differentiation (14) also points towards stochastic selection. This does not entirely rule out selection because the appropriate antigen presenting cell may still be required of γδ maturation.

Not only does a large fraction γδ thymocytes appear to be derived from distinct DN1 thymocytes prior to TCR signaling, but we also observed compartmentalization of effector outcomes. Unlike αβ T cells, γδ T cells are thought to acquire their effector potential in the thymus rather than upon antigen exposure in secondary lymphoid organs (41). Although none of the DN1 subpopulations expressed definitive markers of specific mature T cell populations, we observed substantial transcriptomic overlap between the IL-17-primed DN1 subpopulations with mature IL-17-expressing γδ thymocytes and between the IFNγ-primed DN1 subpopulations with mature IFNγ-expressing γδ thymocytes. Expression of key transcription factors was notable. We showed that DN1d thymocytes express many of transcription factors that are expressed by mature IL-17-producing γδ thymocytes but not IFNγ-producing thymocytes, like *Bcl11b, Etv5, Sox13, Rcf7, Rorc and Maf*. Sox13 has previously be shown to be an important lineage determining factor for the neonatal IL-17A-producing cells (18). It is thought to act in concert with other transcription factors like *Bcl11b, Tcf7, Rorc* and *Maf* to specify the IL-17A-effector program in γδ T cells (18, 42).

While DN1e-1 and DN1e-2 thymocytes were also biased towards a Vγ2^+^ IL-17A-producing γδ T cell fate, only DN1e-3 and DN1e-4 thymocytes displayed the plasticity to also produce Vγ1^+^ IFNγ^+^ cells. This plasticity is likely to involve the differential expression of transcription factors that contribute to distinct effector fates. Although all four DN1e subpopulations expressed *Stat1*, DN1e-1 and DN1e-2 also expressed *Maf*. c-Maf is known to positively regulate IL-17A-producing γδ T cell development (43), whereas a lack of c-Maf expression by γδ T cells correlates with increased IFNγ expression (43-45). Furthermore DN1e-1 cells were found to express *Gata3*, encoding another important regulator of IFNγ expression (46). The interplay between transcription factors may thus be key to lineage decisions. Thus, critical components of the IL-17 or IFNγ γδ T cell transcriptional programs appear to be already in place within distinct DN1 subpopulations, suggesting that predetermination contributes to γδ effector subset differentiation. Further analysis of chromatin states and epigenetic mechanisms associated with these transcription factors will likely be valuable for revealing the regulatory cascades that drive the different γδ effector outcomes.

## Acknowledgments

We thank Lucy Kloboucnik in the St Vincent’s Hospital Melbourne’s BioResources Centre for expert animal husbandry, and Anthony Di Carluccio from St Vincent’s Institute’s Cytometry Facility and Casey Anttila from Walter and Eliza Hall Institute’s Cytometry Facility for assistance with cell sorting and the scRNAseq libraries. We also thank Dr David Izon (St Vincent’s Institute) for the OP9-DL1 cell line, who originally obtained them from Dr Juan Carlos Zúñiga-Pflücker (University of Toronto).

---

**SUPPLEMENTAL FIGURE 1.**
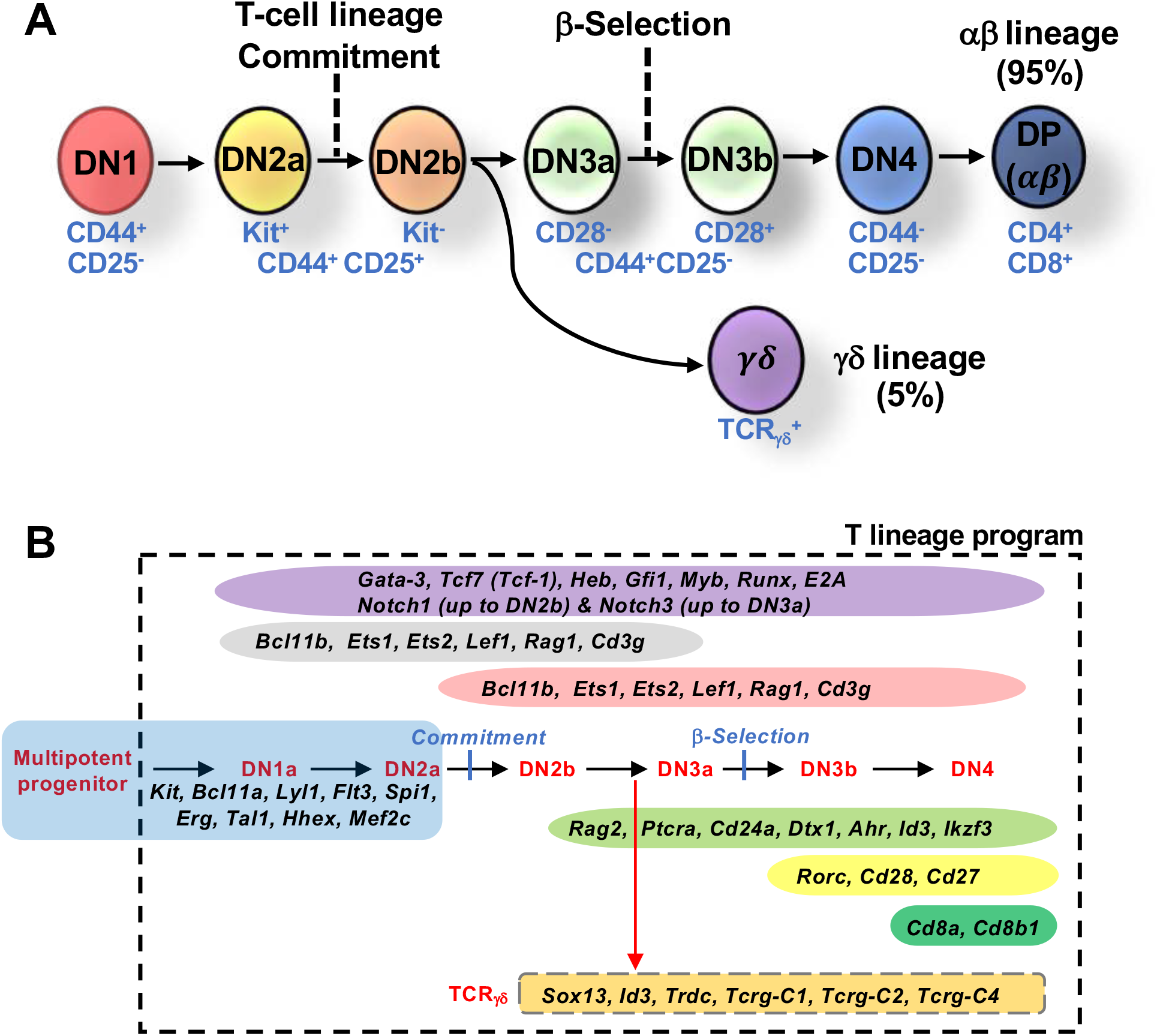
The standard model of murine T cell development. **(A)** T cells are divided into two main lineages, αβ and γδ T cells, which are defined by T cell receptor (TCR) chain expression. Shown is a schematic overview of early T cell development, which is divided into four stages, termed double negative (DN) 1 to 4, based on expression of key cell surface markers. **(B)** The key cell surface markers are indicated. The current model assumes the γδ lineage branches off from αβ lineage during DN2b>DN3a stage, when *Tcrb/g/d* gene rearrangements occur. Also shown is a summary of the expression patterns of key marker genes that define the DN stages of T cell development as defined at a population level.

**SUPPLEMENTAL FIGURE 2.**
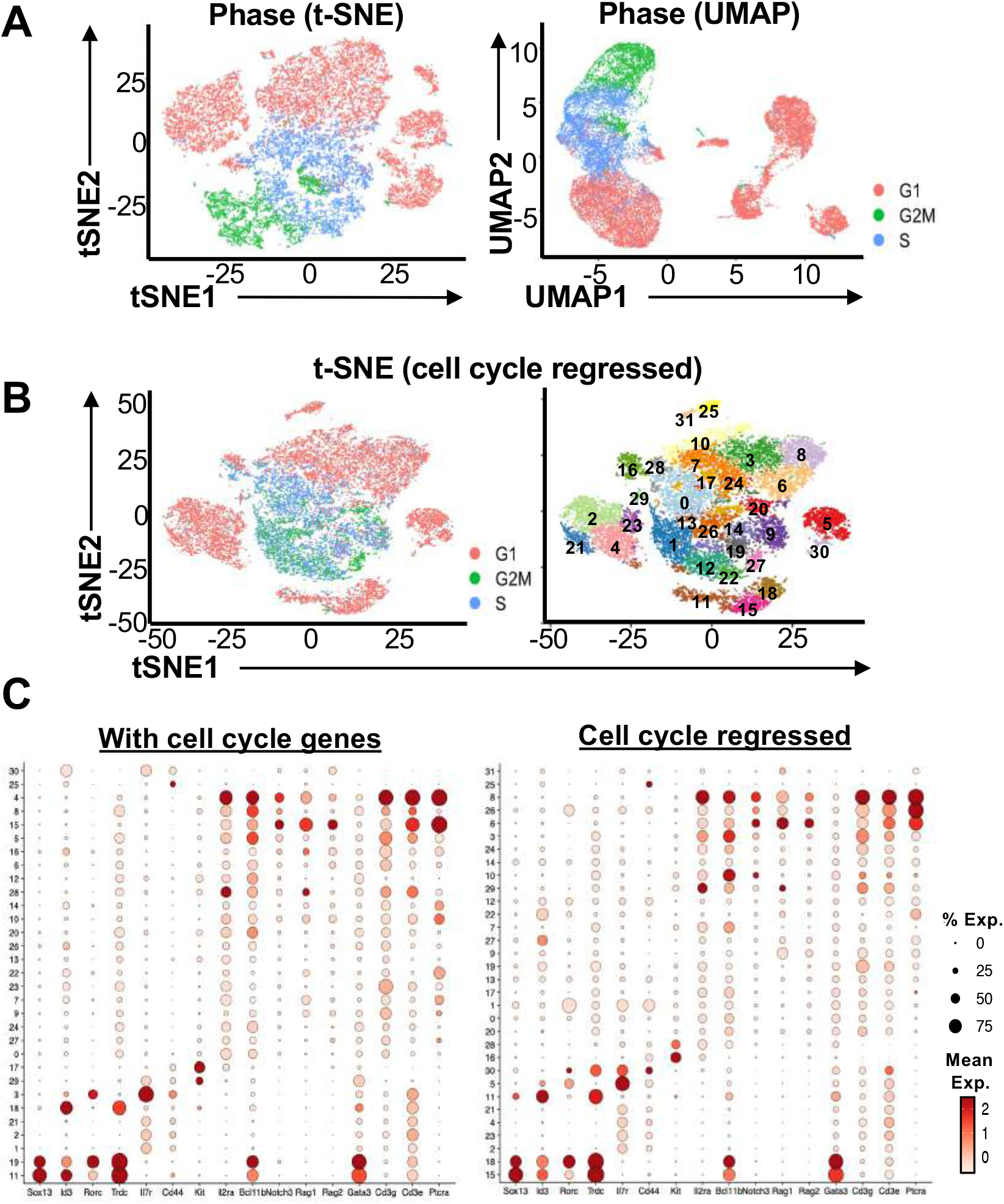
Cell cycle has a minimal impact on the clustering of the DN thymocyte scRNA-seq data. **(A)** t-SNE (left) and UMAP (right) visualization of the integrated scRNA-seq data derived from the three 10X runs of DN and TCRγδ+ thymocytes. Each cell was tagged as in G1, G2/M or S phase based on expression of cell cycle genes. **(B)** Cell cycle genes were first regressed out using Seurat’s built-in regression model and clustered. The cells were then retagged to cell cycle stage (left). 31 distinct clusters were resolved (right). **(C)** Dot plot showing the expression of key markers genes across the clusters comparing the output with cell cycle genes left in or regressed out. Dot size indicates the percentage of cells within the cluster expressing the gene, while color saturation indicates average expression.

**SUPPLEMENTAL FIGURE 3.**
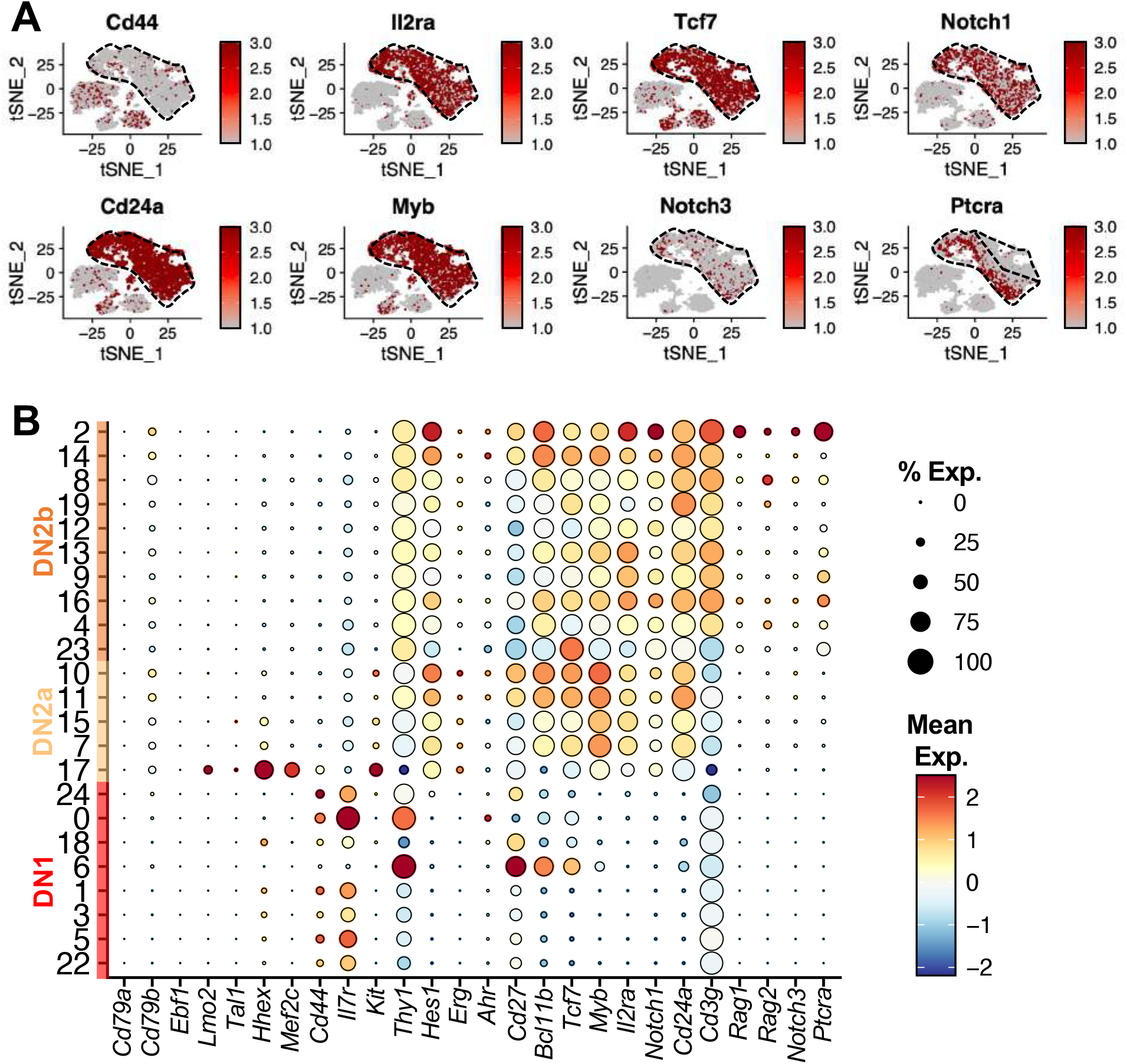
Single-cell analysis of DN1 and DN2 thymocytes. **(A)** Feature plots of some of the markers used to define the clusters in Main Fig. 3A. **(B)** Dot plot showing the expression of key markers genes previously shown to be differentially expressed between DN1, DN2a and DN2b. Cluster 20, 21 and 25 were identified as non-thymocytes (doublets and B cells) and were removed from downstream analyses. Dot size indicates the percentage of cells within each cluster expressing the gene, while color saturation indicates average expression level within the cluster.

