## Supplemental data for "Distinct subpopulations of DN1 thymocytes exhibit preferential γδ T lineage potential"

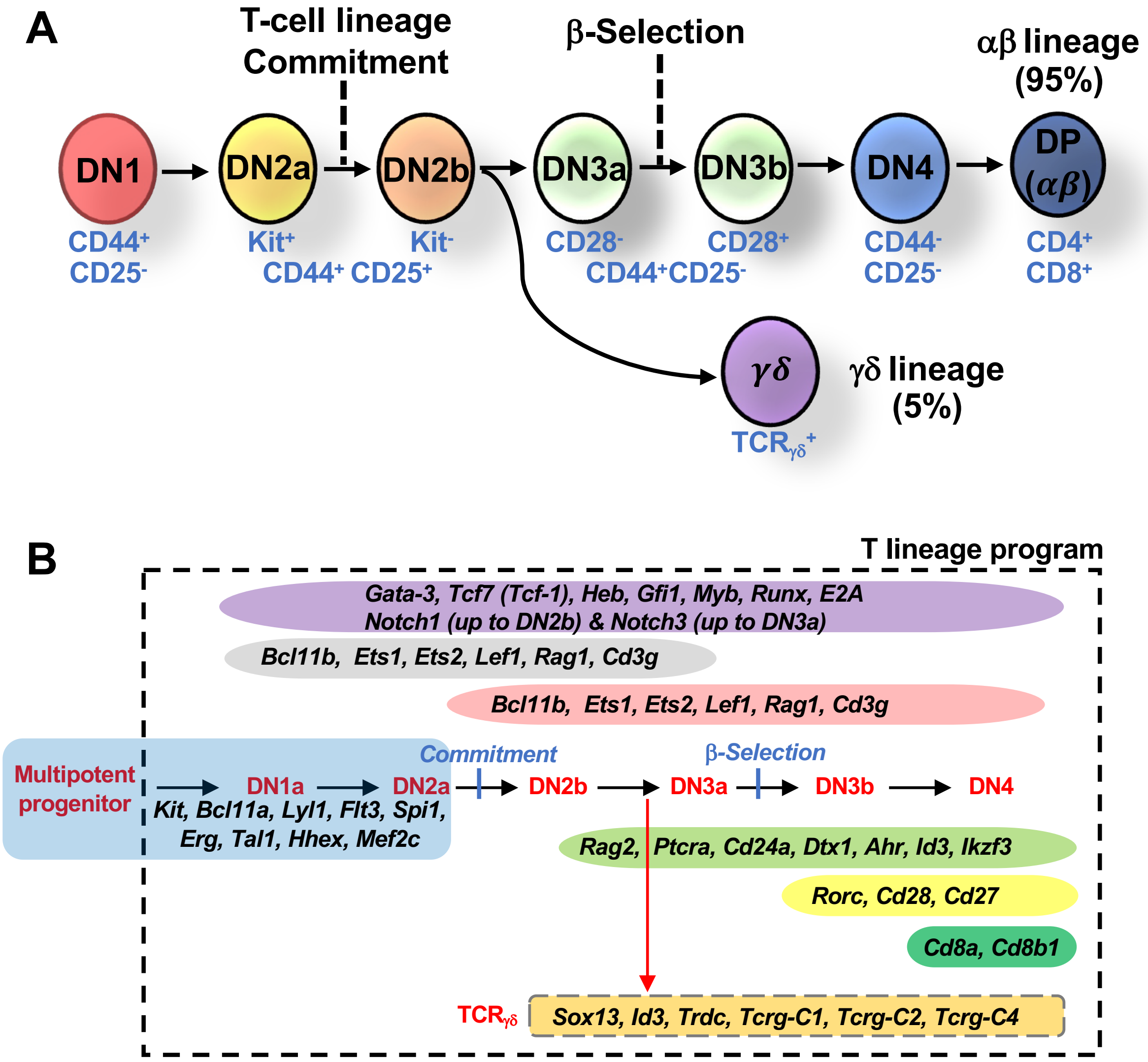


**SUPPLEMENTAL FIGURE 1.** The standard model of murine T cell development. **(A)** T cells are divided into two main lineages, αβ and γδ T cells, which are defined by T cell receptor (TCR) chain expression. Shown is a schematic overview of early T cell development, which is divided into four stages, termed double negative (DN) 1 to 4, based on expression of key cell surface markers. **(B)** The key cell surface markers are indicated. The current model assumes the γδ lineage branches off from αβ lineage during DN2b>DN3a stage, when *Tcrb/g/d* gene rearrangements occur. Also shown is a summary of the expression patterns of key marker genes that define the DN stages of T cell development as defined at a population level.

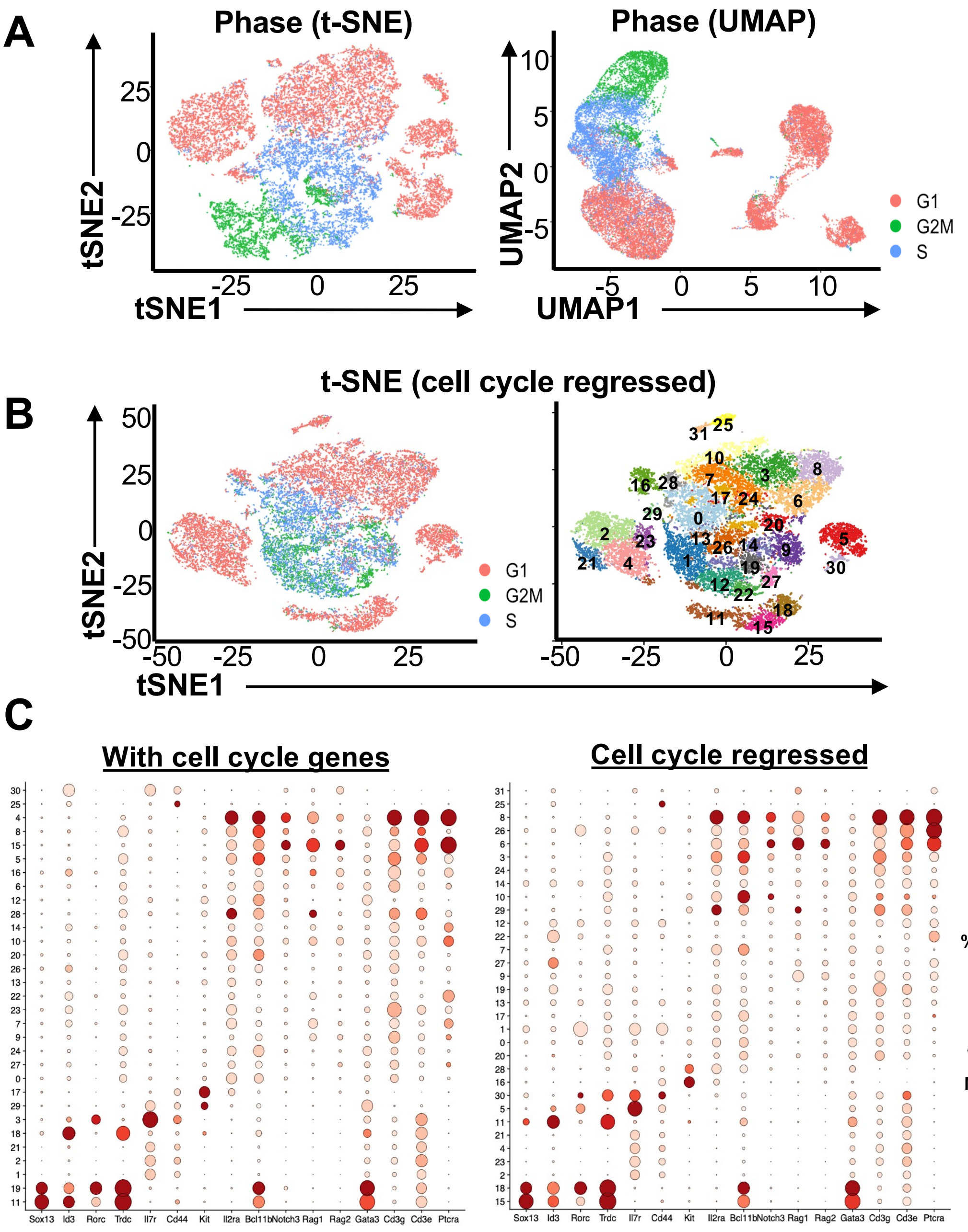


**SUPPLEMENTAL FIGURE 2.** Cell cycle has a minimal impact on the clustering of the DN thymocyte scRNA-seq data. **(A)** t-SNE (left) and UMAP (right) visualization of the integrated scRNA-seq data derived from the three 10X runs of DN and TCRγδ$^+$ thymocytes. Each cell was tagged as in G1, G2/M or S phase based on expression of cell cycle genes. **(B)** Cell cycle genes were first regressed out using Seurat's built-in regression model and clustered. The cells were then retagged to cell cycle stage (left). 31 distinct clusters were resolved (right). **(C)** Dot plot showing the expression of key markers genes across the clusters comparing the output with cell cycle genes left in or regressed out. Dot size indicates the percentage of cells within the cluster expressing the gene, while color saturation indicates average expression.

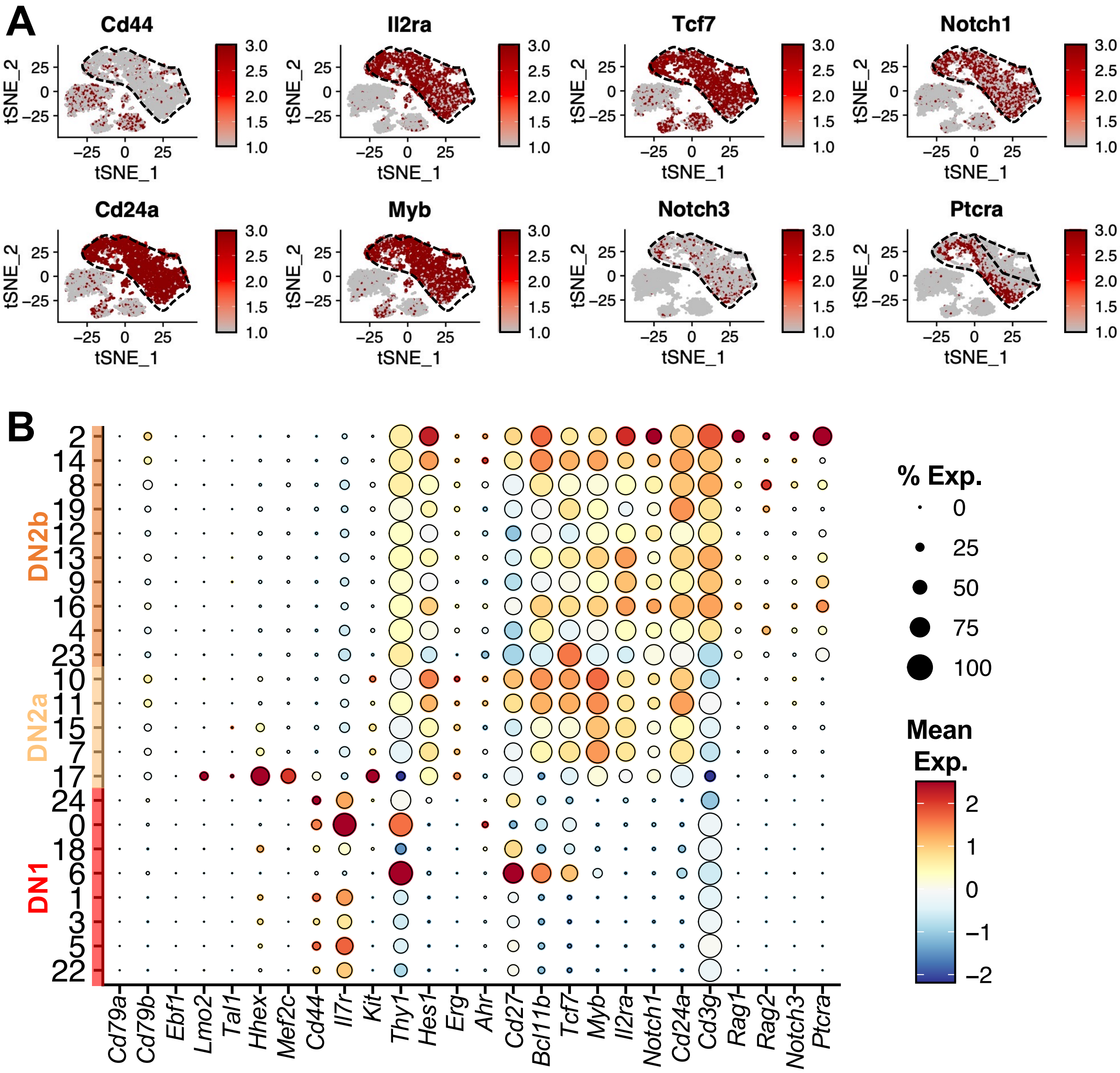


**SUPPLEMENTAL FIGURE 3.** Single-cell analysis of DN1 and DN2 thymocytes. **(A)** Feature plots of some of the markers used to define the clusters in Main Fig. 3A. **(B)** Dot plot showing the expression of key markers genes previously shown to be differentially expressed between DN1, DN2a and DN2b. Cluster 20, 21 and 25 were identified as non-thymocytes (doublets and B cells) and were removed from downstream analyses. Dot size indicates the percentage of cells within each cluster expressing the gene, while color saturation indicates average expression level within the cluster.

**SUPPLEMENTAL TABLE 1.** *P*-values comparing αβ versus γδ differentiation from DN1 subpopulations after 14d on OP9-DL1 monolayer (related to Figure 4B)

| Comparing % $CD8^+$ | | | | | | | | | |
|---|---|---|---|---|---|---|---|---|---|
| ***p-value*** | **DN1** | **1a** | **1b** | **1c** | **1d** | **1e-1** | **1e-2** | **1e-3** | **1e-4** |
| **DN1** | | ns | **** | **** | **** | **** | **** | **** | **** |
| **1a** | ns | | **** | **** | **** | **** | **** | **** | **** |
| **1b** | **** | **** | | ns | **** | **** | **** | **** | **** |
| **1c** | **** | **** | ns | | **** | **** | **** | **** | **** |
| **1d** | **** | **** | **** | **** | | ns | ns | ns | ns |
| **1e-1** | **** | **** | **** | **** | ns | | ns | ns | ns |
| **1e-2** | **** | **** | **** | **** | ns | ns | | ns | ns |
| **1e-3** | **** | **** | **** | **** | ns | ns | ns | | ns |
| **1e-4** | **** | **** | **** | **** | ns | ns | ns | ns | |
| **Comparing % TCRγδ$^+$** | | | | | | | | | |
| ***p-value*** | **DN1** | **1a** | **1b** | **1c** | **1d** | **1e-1** | **1e-2** | **1e-3** | **1e-4** |
| **DN1** | | ns | ns | ns | ns | **** | *** | * | **** |
| **1a** | ns | | ns | ns | * | **** | **** | **** | **** |
| **1b** | ns | ns | | ns | * | **** | **** | **** | **** |
| **1c** | ns | ns | ns | | ns | **** | **** | **** | **** |
| **1d** | ns | * | * | ns | | ** | ns | ns | ns |
| **1e-1** | **** | **** | **** | **** | ** | | ns | * | ns |
| **1e-2** | **** | **** | **** | **** | ns | ns | | ns | ns |
| **1e-3** | * | **** | **** | **** | ns | * | ns | | ns |
| **1e-4** | *** | **** | **** | **** | ns | ns | ns | ns | |

The data was analyzed by one-way analysis of variance (ANOVA). ns = not significant, *$P<0.05$, ** $P<0.01$, *** $P<0.001$, **** $P<0.0001$.

**SUPPLEMENTAL TABLE 2.** *P*-values comparing αβ versus γδ differentiation from DN1 subpopulations after 20d on OP9-DL1 monolayer (related to Figure 4B)

| Comparing % CD8$^+$ | | | | | | | | | |
|---|---|---|---|---|---|---|---|---|---|
| ***p-value*** | **DN1** | **1a** | **1b** | **1c** | **1d** | **1e-1** | **1e-2** | **1e-3** | **1e-4** |
| **DN1** | | ns | ns | **** | **** | **** | **** | **** | **** |
| **1a** | ns | | ns | **** | **** | **** | **** | **** | **** |
| **1b** | ns | ns | | **** | **** | **** | **** | **** | **** |
| **1c** | **** | **** | **** | | **** | **** | **** | **** | **** |
| **1d** | **** | **** | **** | **** | | ns | ns | ns | ns |
| **1e-1** | **** | **** | **** | **** | ns | | ns | ns | ns |
| **1e-2** | **** | **** | **** | **** | ns | ns | | ns | ns |
| **1e-3** | **** | **** | **** | **** | ns | ns | ns | | ns |
| **1e-4** | **** | **** | **** | **** | ns | ns | ns | ns | |
| **Comparing % TCRγδ$^+$** | | | | | | | | | |
| ***p-value*** | **DN1** | **1a** | **1b** | **1c** | **1d** | **1e-1** | **1e-2** | **1e-3** | **1e-4** |
| **DN1** | | ns | ns | ns | ns | **** | **** | * | *** |
| **1a** | ns | | ns | ns | ** | **** | **** | *** | **** |
| **1b** | ns | ns | | ns | *** | **** | **** | **** | **** |
| **1c** | ns | ns | ns | | ns | **** | **** | ns | * |
| **1d** | ns | ** | *** | ns | | * | ** | ns | ns |
| **1e-1** | **** | **** | **** | **** | * | | ns | ** | ns |
| **1e-2** | **** | **** | **** | **** | ** | ns | | *** | * |
| **1e-3** | * | *** | **** | ns | ns | ** | *** | | ns |
| **1e-4** | *** | **** | **** | * | ns | ns | * | ns | |

The data was analyzed by one-way analysis of variance (ANOVA). ns = not significant, * $P<0.05$, ** $P<0.01$, *** $P<0.001$, **** $P<0.0001$.

**SUPPLEMENTAL TABLE 3.** *P*-values comparing αβ versus γδ differentiation from DN2 subpopulations after 14d on OP9-DL1 monolayer (related to Figure 4D)

**Comparing % $CD8^+$**

| *p-value* | DN2 | 2a-1 | 2a-2 | 2a-3 | 2a-4 | 2b-1 | 2b-2 | 2b-3 | 2b-4 | 2b-5 | 2b-6 | 2b-7 |
|---|---|---|---|---|---|---|---|---|---|---|---|---|
| **DN2** | | * | **** | *** | **** | ns | ns | *** | ns | ns | ns | ns |
| **2a-1** | * | | ns | ns | ns | **** | ns | **** | *** | ** | ns | **** |
| **2a-2** | **** | ns | | ns | ns | **** | *** | **** | **** | **** | ns | **** |
| **2a-3** | *** | ns | ns | | ns | **** | *** | **** | **** | **** | ns | **** |
| **2a-4** | **** | ns | ns | ns | | **** | **** | **** | **** | **** | ns | **** |
| **2b-1** | ns | **** | **** | **** | **** | | * | ns | ns | ns | **** | ns |
| **2b-2** | ns | ns | *** | *** | **** | * | | **** | ns | ns | ns | ns |
| **2b-3** | *** | **** | **** | **** | **** | ns | **** | | * | *** | **** | ns |
| **2b-4** | ns | *** | **** | **** | **** | ns | ns | * | | ns | ** | ns |
| **2b-5** | ns | *** | **** | **** | **** | ns | ns | *** | ns | | * | ns |
| **2b-6** | ns | ns | ns | ns | ns | **** | ns | **** | ** | * | | *** |
| **2b-7** | ns | **** | **** | **** | **** | ns | ns | ns | ns | ns | *** | |

**Comparing % $TCR\gamma\delta^+$**

| *p-value* | DN2 | 2a-1 | 2a-2 | 2a-3 | 2a-4 | 2b-1 | 2b-2 | 2b-3 | 2b-4 | 2b-5 | 2b-6 | 2b-7 |
|---|---|---|---|---|---|---|---|---|---|---|---|---|
| **DN2** | | ns | * | * | ns | ns | ns | ns | ns | ns | ns | ns |
| **2a-1** | ns | | * | * | ns | ns | ns | ns | ns | ns | ns | ns |
| **2a-2** | * | * | | ns | ns | ** | * | *** | ns | ns | ns | * |
| **2a-3** | * | * | ns | | ns | * | ns | ** | ns | ns | ns | ns |
| **2a-4** | ns | ns | ns | ns | | ns | ns | * | ns | ns | ns | ns |
| **2b-1** | ns | ns | ** | * | ns | | ns | ns | ns | ns | ns | ns |
| **2b-2** | ns | ns | * | ns | ns | ns | | ns | ns | ns | ns | ns |
| **2b-3** | ns | ns | *** | ** | * | ns | ns | | ns | ns | ns | ns |
| **2b-4** | ns | ns | ns | ns | ns | ns | ns | ns | | ns | ns | ns |
| **2b-5** | ns | ns | ns | ns | ns | ns | ns | ns | ns | | ns | ns |
| **2b-6** | ns | ns | ns | ns | ns | ns | ns | ns | ns | ns | | ns |
| **2b-7** | ns | ns | * | ns | ns | ns | ns | ns | ns | ns | ns | |

The data was analyzed by one-way analysis of variance (ANOVA). ns = not significant, * $P<0.05$, ** $P<0.01$, *** $P<0.001$, **** $P<0.0001$.

**SUPPLEMENTAL TABLE 4.** *P*-values comparing αβ versus γδ differentiation from DN2 subpopulations after 20d on OP9-DL1 monolayer (related to Figure 4D)

| **Comparing % $CD8^+$** | | | | | | | | | | | | |
|---|---|---|---|---|---|---|---|---|---|---|---|---|
| ***p-value*** | **DN2** | **2a-1** | **2a-2** | **2a-3** | **2a-4** | **2b-1** | **2b-2** | **2b-3** | **2b-4** | **2b-5** | **2b-6** | **2b-7** |
| **DN2** | | ns | ns | ns | ns | ns | ns | ns | * | * | ns | ns |
| **2a-1** | ns | | ns | ns | ns | ns | ns | ns | ** | ** | ns | ns |
| **2a-2** | ns | ns | | ns | ns | ns | ns | ns | ns | ns | ns | ns |
| **2a-3** | ns | ns | ns | | ns | ns | ns | ns | ns | ns | ns | ns |
| **2a-4** | ns | ns | ns | ns | | ns | ns | ns | ns | ns | ns | ns |
| **2b-1** | ns | ns | ns | ns | ns | | ns | ns | ** | ** | ns | ns |
| **2b-2** | ns | ns | ns | ns | ns | ns | | ns | ns | ns | ns | ns |
| **2b-3** | ns | ns | ns | ns | ns | ns | ns | | ns | ns | ns | ns |
| **2b-4** | * | ** | ns | ns | ns | ** | ns | ns | | ns | ns | ** |
| **2b-5** | * | ** | ns | ns | ns | ** | ns | ns | ns | | ns | *** |
| **2b-6** | ns | ns | ns | ns | ns | ns | ns | ns | ns | ns | | ns |
| **2b-7** | ns | ns | ns | ns | ns | ns | ns | ns | ** | *** | ns | |
| **Comparing % TCRγδ$^+$** | | | | | | | | | | | | |
| ***p-value*** | **DN2** | **2a-1** | **2a-2** | **2a-3** | **2a-4** | **2b-1** | **2b-2** | **2b-3** | **2b-4** | **2b-5** | **2b-6** | **2b-7** |
| **DN2** | | ns | ns | ns | ns | ns | ns | ns | ns | ns | ns | ns |
| **2a-1** | ns | | ** | ns | ns | ns | ns | ns | ** | ns | ns | ns |
| **2a-2** | ns | ** | | ** | ns | ns | ns | ns | ns | ** | * | ns |
| **2a-3** | ns | ns | ns | | ns | ns | ns | ns | ** | ns | ns | ns |
| **2a-4** | ns | ns | ns | ns | | ns | ns | ns | * | ns | ns | ns |
| **2b-1** | ns | ns | ns | ns | ns | | ns | ns | ns | ns | ns | ns |
| **2b-2** | ns | ns | ns | ns | ns | ns | | ns | * | ns | ns | ns |
| **2b-3** | ns | ns | ns | ns | ns | ns | ns | | ns | ns | ns | ns |
| **2b-4** | ns | ** | ns | ** | * | ns | * | ns | | ** | * | ns |
| **2b-5** | ns | ns | ** | ns | ns | ns | ns | ns | ** | | ns | ns |
| **2b-6** | ns | ns | * | ns | ns | ns | ns | ns | * | ns | | ns |
| **2b-7** | ns | ns | ns | ns | ns | ns | ns | ns | ns | ns | ns | |

The data was analyzed by one-way analysis of variance (ANOVA). ns = not significant, * $P<0.05$, ** $P<0.01$, *** $P<0.001$, **** $P<0.0001$.

**SUPPLEMENTAL TABLE 5.** *P*-values comparing $V\gamma1^+$ TCRγδ versus $V\gamma2^+$ TCRγδ cells produced from DN1 subpopulations after 14d on OP9-DL1 monolayer (related to Figure 4D)

| **Comparing % $V\gamma1^+$ TCRγδ** | | | | | | | |
|---|---|---|---|---|---|---|---|
| ***p-value*** | **DN1** | **1c** | **1d** | **1e-1** | **1e-2** | **1e-3** | **1e-4** |
| **DN1** | | ns | ns | **** | *** | ns | ns |
| **1c** | ns | | * | **** | **** | ns | * |
| **1d** | ns | * | | * | * | ns | ns |
| **1e-1** | **** | **** | * | | ns | **** | ** |
| **1e-2** | *** | **** | * | ns | | *** | ** |
| **1e-3** | ns | ns | ns | **** | *** | | ns |
| **1e-4** | ns | * | ns | ** | ** | ns | |
| **Comparing % $V\gamma2^+$ TCRγδ** | | | | | | | |
| ***p-value*** | **DN1** | **1c** | **1d** | **1e-1** | **1e-2** | **1e-3** | **1e-4** |
| **DN1** | | ns | ns | ns | ns | ns | ns |
| **1c** | ns | | ns | ns | ns | ns | ns |
| **1d** | ns | ns | | ns | ns | ns | ns |
| **1e-1** | ns | ns | ns | | ns | ns | ns |
| **1e-2** | ns | ns | ns | ns | | ns | ns |
| **1e-3** | ns | ns | ns | ns | ns | | ns |
| **1e-4** | ns | ns | ns | ns | ns | ns | |

The data was analyzed by one-way analysis of variance (ANOVA). ns = not significant, * $P<0.05$, ** $P<0.01$, *** $P<0.001$, **** $P<0.0001$.
